# E2EDA: Protein domain assembly based on end-to-end deep learning

**DOI:** 10.1101/2023.01.25.525481

**Authors:** Hai-Tao Zhu, Yu-Hao Xia, Guijun Zhang

## Abstract

With the development of deep learning, almost all single-domain proteins can be predicted at experimental resolution. However, the structure prediction of multi-domain proteins remains a challenge. Achieving end-to-end protein domain assembly and further improving the accuracy of the full-chain modelling by accurately predicting inter-domain orientation while improving the assembly efficiency will provide significant insights into structure-based drug discovery. In addition, the available GPU memory limits the size of a full-chain protein which can be predicted. Based on the divide-and-conquer strategy, the single-domain structure is predicted by the state-of-the-art prediction method, such as AlphaFold2, and then assembled into a full-chain model through the domain assembly method, which can effectively reduce the demand for hardware resources. In this work, we propose an End-To-End Domain Assembly method based on deep learning, named E2EDA. We first develop an EffificientNetV2-based deep learning model (RMNet), which is specialised for predicting inter-domain orientations. The RMNet uses an attention mechanism to predict inter-domain rigid motion by fusing sequence features, multiple template features and single-domain features. Then, the predicted rigid motions are converted into inter-domain spatial transformations to assemble full-chain models of multi-domain proteins directly without time-consuming simulation processes. Finally, a scoring strategy, RMscore, is designed to select the best model from multiple assembled models to improve assembly accuracy. The experimental results show that the average TM-score of the model assembled by E2EDA on the benchmark set (356) is 0.84, which is better than other domain assembly methods SADA (0.80), DEMO (0.74) and AIDA (0.63). Meanwhile, on our constructed human protein dataset from AlphaFold DB, the model reassembled by E2EDA is 6.8% higher than the full-chain model predicted by AlphaFold2, indicating that E2EDA can capture more accurate inter-domain orientations to improve the quality of the model predicted by AlphaFold2. Furthermore, the average running time of E2EDA on the benchmark is reduced by 74.6% compared with the domain assembly simulation method SADA, which indicates that E2EDA can effectively improve assembly efficiency through an end-to-end manner.The online server is at http://zhanglab-bioinf.com/E2EDA/.

## 1 Introduction

Domains are typically defined either as ‘folding units’ using structural definitions or as ‘independent evolutionary units’ using evolution-based definitions. Although the two definitions are distinct, the resulting family of domains is equivalent in most cases, suggesting that domains are the fundamental units of protein structure and evolution [1]. Proteins are the expression products of genes; genes generate new genes through recombination, replication, fission and fusion. Proteins are also formed by replicating and combining in different ways through a limited family of domains [2]. Each protein domain is folded into a compact three-dimensional structure that is conserved among different multi-domain proteins [3]. Two-thirds of the structures in the protein database are single-domain proteins [4]. However, most proteins in nature contain multiple domains, which indicates that the current experimental methods have difficulty determining the structure of multi-domain proteins. Meanwhile, with the advancement of technology in single-domain protein structure prediction, determining the structure of multi-domain proteins by computational methods has become increasingly important.

The structure prediction of multi-domain proteins has two typical methods. One is the simulation-based domain assembly method, and the other is the full-chain modelling method based on deep learning. Simulation-based domain assembly methods generally have two ways to assemble multi-domain protein structures from single-domain structures. The first way of domain assembly is to sample the degrees of freedom of the linker. In this method, the domain assembly can be viewed as a problem of *de novo* prediction of relatively short amino acid sequences with preformed N- and C-terminal structures [5]. AIDA [6] is a linker-based domain assembly method, in which the four residues around the domain boundary are regarded as a linker. The degree of freedom of the linker is continuously adjusted through energy minimization simulation, and finally the model with the smallest energy is generated. Furthermore, the domain assembly can also be viewed as a docking problem. In our previous works, DEMO [7] and SADA [8] assembled the single-domain structure by rigid body docking. During domain assembly, SADA and DEMO calculate the average coordinates of the backbone atoms as the centroid of the single-domain structure and guide the sampling of the rigid body degrees of freedom between the two domains by reducing the energy potential. Different from AIDA, DEMO and SADA also used structural analogues as templates during the assembly process. In addition, SADA utilises the distance information predicted by deep learning and constructs it as the energy term of the force field for domain assembly. However, given the imperfection of the energy force field in the traditional domain assembly method, the assembled full-chain model does not always correspond to the native structure. Moreover, the simulation-based domain assembly method is accompanied by a time-consuming assembly simulation process. Therefore, an end-to-end domain assembly method that directly predicts inter-domain orientations through deep learning may help improve the accuracy and efficiency of assembly.

In addition to protein domain assembly methods, full-chain modelling based on deep learning is also one of the mainstream methods for predicting multi-domain proteins. Since CASP11, deep learning techniques have had a major impact on the field of protein structure prediction [9]. From the initial methods of predicting contact, such as RaptorX [10], DMP [11], etc., to the later methods of predicting distance, such as AlphaFold [12], trRosetta [13, 14], etc., have been developed successively. The contact and distance between the residues based on deep learning predictions are used to build energy potential to improve the quality of the predicted protein structure [10, 15-17]. Recently, with the advent of end-to-end approaches such as AlphaFold2 [18] and RoseTTAFold [19], processes no longer rely on traditional energy potentials to guide folded protein structures. The results of Alphafold2 in CASP14 show that compared with the protein structure prediction guided by the energy potential, the end-to-end structure prediction method may be more accurate. For many single-domain proteins, the model predicted by AlphaFold2 can achieve experimental accuracy. However, for multi-domain proteins, as the training set of AlphaFold2 is heavily biased towards single-domain proteins, two-thirds of the training set PDB database is single-domain proteins, which makes AlphaFold2 relatively poor in capturing inter-domain orientations during full-chain modelling [20]. In addition, for large proteins (e.g. >1,300 amino acids), most available GPU memory may have difficulty meeting the requirements of its inference [18]. Therefore, constructing a deep learning model to predict inter-domain orientations, and based on a divide-and-conquer strategy to first predict single-domain structures and then assemble them into a full-chain model, may help improve accuracy while reducing the demand for hardware. In addition, in the modelling of multi-domain proteins, attention is paid to improving the quality of MSA while ignoring template-based modelling methods. Breakthroughs in single-domain modelling, such as AlphaFold2, provide a possibility for template-based domain assembly methods.

In this study, we take full advantage of advances in single-domain prediction techniques and further consider the prediction of the inter-domain orientation of multi-domain proteins. Therefore, we propose E2EDA, an end-to-end domain assembly method based on deep learning. E2EDA predicts rigid motions between domains, which are translated into inter-domain spatial transformations to assemble full-chain models directly without time-consuming simulations. The experimental results show that, on the benchmark dataset and the human protein dataset, E2EDA improves the modelling accuracy of multi-domain proteins and significantly improves the assembly efficiency.

## 2 Materials and Methods

As illustrated in Figure 1A, E2EDA is an end-to-end protein domain assembly method based on deep learning. The pipeline consists of two main steps: inter-residue rigid motion prediction and domain assembly using rigid motion. For a given sequence, the corresponding MSA and template are generated by HHblits and HHsearch [21]. Then, the RMNet uses the attention mechanism to fuse sequence features, single-domain features and multiple template features to predict rigid motion distribution, which contains six components consisting of quaternions representing rotation and vectors representing translation. In the assembly stage, multiple rigid motions with the highest confidence among inter-domain residues are selected from the predicted rigid motion distribution to assemble the multi-domain protein. Finally, a scoring function is constructed from the predicted rigid motion distribution to score the multi-domain proteins assembled using rigid motion. The five models with the best scores are selected as the final model.

**Figure. 1.**
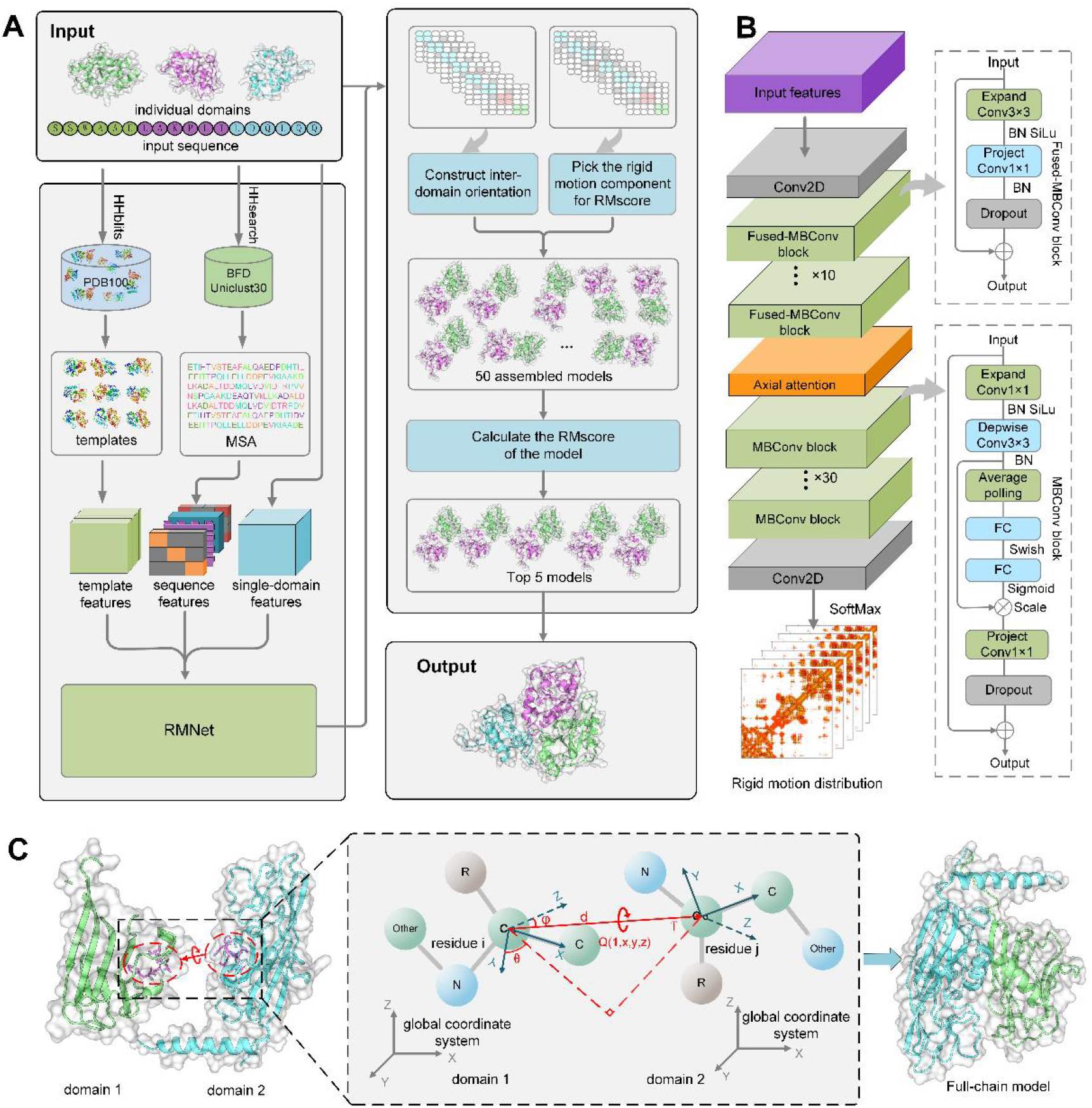
End-to-end protein domain assembly by predicting inter-domain rigid motion. (A) The pipeline of E2EDA. The proposed E2EDA first predicts the rigid motion between residues by the prediction network RMNet. Then, multiple models are directly assembled using the predicted rigid motions without domain assembly simulations. The final model is selected by scoring the assembled model through RMscore. (B) Architecture of RMNet based on EffificientNetV2 network. (C) Assembly of multi-domain proteins using predicted inter-domain rigid motions.

### 2.1 Rigid motion prediction

In this work, we predict rigid motions that adequately describe the relative positional relationship of C_α_ atoms between two residues. As shown in Figure 1C, we denote rigid motion by (*M* = *Q, T*), where *Q* ∈ *R*^4^ represents the quaternion used to construct the rotation matrix between domains. *T* ∈ *R*^3^ is used to construct the translation vector between domains. For each residue, we take the C_α_ atom as the coordinate origin. Then, we construct a local coordinate system using three atoms: N, C_α_ and C. Next, we calculate the rigid motion *M*_*i*_ from this coordinate system to the global coordinate system. The detailed procedure for constructing *M*_*i*_ can be found in Supplementary Method S1. The rigid motion *M*_*ij*_ between the *i*-th residue and the *j*-th residue can be derived from *M*_*i*_ and *M*_*j*_, as shown in Equation (1) [18]. The detailed calculation process can be found in Supplementary Method S2.

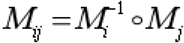

where 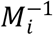 denotes the inverse of *M*_*i*_ and the ° operator denotes a combination of rigid motions. The network RMNet predicts six scalars, namely, *x, y, z, d, φ* and *θ*, which together constitute the rigid motion between residues. *x, y* and *z* are constructed as the imaginary part of *Q*, and *d, φ* and *θ* are used to construct the translation.

#### 2.1.1 Datasets

To train a neural network to predict rigid motion, we construct a non-redundant dataset of 9,803 multi-domain proteins. Firstly, we cluster the in-housed multi-domain database MPDB (As of June 2022, 48,224 multi-domain proteins are available) [8] using CD-HIT [22] with a 40% sequence identity cutoff, resulting in 10,593 multi-domain proteins. Then, we use DomainParser [23] to decompose these multi-domain proteins, and the proteins that cannot be decomposed are removed. Thus, 9,803 proteins are left, of which 2,985 proteins have more than 425 residues. In addition, 1,000 of 2,985 large proteins (>425 residues) are randomly selected to be added to the training set at each iteration of training. The final dataset contains 9,803 proteins, 95% of which are used for training and 5% for validation.

#### 2.1.2 Feature design

As shown in Figure 1A, the input features of RMNet are mainly composed of three parts: MSA features, homologous template features and single-domain features. MSA contains co-evolutionary information of target proteins, and recent studies have shown that the accuracy of network prediction depends on the quality of the input MSA [14]. In this work, in addition to the sequence database Uniclust30 [24], HHblits also searches the BFD database containing metagenomics, metatranscriptomes and reference databases to generate higher-quality MSA [18]. The e-value cutoff of HHblits is gradually relaxed according to 1e-30, 1e-10, 1e-6 and 1e-3 until the homologous sequences with 75% coverage in MSA are more than 2,000 or the homologous sequences with 50% coverage are more than 4,000. If the collected sequences are still not enough, we directly use the e-value of 1e-3 to search for MSA. The MSA features are calculated and extracted from the MSA in real-time, which has a total of 526 feature channels. The sequence features include: one-hot encoding of the input sequence (*L* × 20, where *L* represents the length of the input sequence), location specificity frequency matrix (*L* × 21, where 20 amino acid types and 1 gap) and positional entropy (*L* × 1). These features are stacked horizontally and vertically to form a 42 × 2 feature map. In addition, residual pair features extracted from MSA include coupling matrix (*L* × *L* × 442) and mean product correction (*L* × *L* × 1).

As the structure of single-domain proteins is known at the time of domain assembly, we consider extracting features from single-domain proteins to improve prediction accuracy. Domains are the fundamental units of protein structure and evolution [1] and proteins are also formed by duplicating and combining in different ways through limited domain families [2], which indicates that the combination of different domains may contain evolutionary information of multi-domain proteins. For each single-domain structure, we construct a local coordinate system using Cα, C and N of each residue. Then, we calculate the rigid motion *M*_*i*_ from the local coordinate system of each residue to the global coordinate system and then convert it into a rigid motion *M*_*ij*_ between residue pairs. Finally, all single-domain features are concatenated into a feature map (*L* × *L* × 6). In addition, we also one-hot encode the boundary information of each domain into a feature map (*L* × *L* × 9). A total of 15 channels for single-domain features exist.

Traditional template-based modelling has certain advantages over other structure prediction methods because homologous templates contain similar structural information [25]. Therefore, template-based features may improve the accuracy of network predictions. In this work, we detect homologous templates of target sequences from PDB100 (before May 03, 2021) by HHsearch. A template is considered a good template when the probability of the template and query sequence is greater than 60% or the e-value is less than 0.001. Template features are extracted from the top 10 good templates; otherwise, if the number of good templates is less than 10 or none, all templates or none are used. The rigid motion between the residues of each template is calculated as the feature of the template. The features of all templates are aggregated together as the final template features (*N* × *L* × *L* × 6). Details of all the features used by E2EDA are listed in Supplementary Table S1.

#### 2.1.3 Network architecture and training

Unlike other methods of predicting distance and contact, we predict rigid motion, which includes six components, namely, *x, y, z, d, φ* and *θ*, as shown in Figure 1C. The overall architecture of the network is shown in Figure 1B. Different from existing ResNet-based methods, a recently proposed network architecture EffificientNetV2 [26] is used, aiming to use resources more efficiently. The EffificientNetV2 architecture can train larger proteins under the same hardware resources, which is more suitable for training multi-domain proteins. After the first layer (2D convolution with filter size 1), the channels of MSA and single-domain features (*L* × *L* × 541) are transformed to 64 and fed into 10 Fused-MBConv blocks. Multiple template features (*N*× *L* × *L* × 6) are embedded via axial attention, which are then fed into a stack of 30 MBConv blocks. The detailed configuration of each block is shown in Supplementary Table S2. After the last MBConv block, the network branches into six independent paths, one for each objective, where each path consists of a 2D convolution and SoftMax activation. The range of *d* (2 to 20 Å) is binned into 36 equidistant segments, plus one bin for those not in the range. Similarly, the *x, y* and *z* range (−1, 1) are all binned into 24 (+ one bin, when *d* is not in 2 to 20 Å); *φ* and *θ* are binned into 24 and 12, respectively, with 15^°^ segments (+ one bin, when *d* is not in 2 to 20 Å).

During training, we randomly select a subset of sequences from each original MSA for training, which ensures that we run on different inputs each time. Due to the limited GPU memory, only a sequence of fewer than 425 residues can be directly accepted; otherwise, it is cut into contiguous subsequences of 425 residues as input. Meanwhile, to reduce the loss of large protein information, two subsequences were randomly sampled: one from the first half of the original sequence, and the other from the second half. Then, they are spliced together to form a new sequence, as shown in Supplementary Figure S1. We use cross-entropy to measure the loss of the six branches, the sum of which is the total loss. We randomly select 95% of the entire data set as the training set and the remaining 5% as the verification set to train for 100 epochs.

### 2.2 Protein domain assembly

The assembly process of E2EDA is shown in Figure 1A. Firstly, multiple models are assembled using the rigid motion predicted by RMNet. Then, these models are scored using RMscore to select the top five models. The above process is iteratively executed when the number of domains is greater than 2. Two domains are assembled each time. Next, the assembled domain is treated as a new single-domain structure, and it is assembled with other single-domain structures. The detailed process of the protein domain assembly is given in Supplementary Figure S2.

#### 2.2.1 Domain assembly using rigid motion

Domain assembly simulations guided by physics-based and knowledge-based energy potential essentially explore inter-domain orientations to assemble multiple single-domain structures into a full-chain model. Here, we directly use the predicted rigid motions for domain assembly, avoiding the simulation process of exploring orientations between domains and greatly improving the assembly efficiency.

Network RMNet predicts the six components, namely, *x, y, z, d, φ* and *θ* of the rigid motion between residues, where *x, y* and *z* are constructed as the imaginary part of the quaternion *Q. d, φ* and *θ* are constructed as translation vectors. The real part of the predicted non-unit quaternion *Q* is fixed at 1. To ensure that the single-domain structure will not be stretched and deformed during assembly, the non-unit quaternion must be normalised. The process of normalising the non-unit quaternions is as follows:

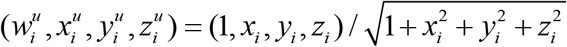

where *x*_*i*_, *y*_*i*_, and *z*_*i*_ are the imaginary part of the non-unit quaternion of residue *i* predicted by the network; 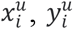 and 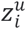 are the imaginary part of the normalised unit quaternion, and 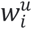 is the real part of the quaternion. A detailed procedure for constructing inter-domain rotation matrices using quaternions for rigid motion can be found in Supplementary Method S3. In addition, *d, φ* and *θ* are used to construct translation vectors between domains. The translation vector is calculated by:

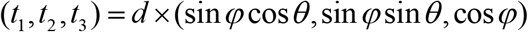

During domain assembly, the lowest confidence of the six components of rigid motion is used as the confidence of the residue pair to ensure that the domain assembly effect will not be too poor due to the inaccurate prediction of a certain component. In addition, if we only use the rigid motion with the highest confidence for domain assembly, the assembled model accuracy may not be the best because the confidence and the accuracy of the predicted rigid motion do not exactly correspond. Therefore, the top 50 rigid motions are selected on the basis of confidence to assemble 50 models. Firstly, according to the selected rigid motion *M*_*ij*_ between the *i*-th residue and the *j*-th residue, *M*_*i*_ and *M*_*j*_ are constructed in the two single-domain structures, respectively. Then, the atomic coordinates of the two single-domain structures are transformed to the corresponding local coordinate systems by PyRosetta [27]. Finally, the atomic coordinates in the residue *j* coordinate system are converted to the residue *i* coordinate system. They are then assembled into a full-chain model, which can be calculated as follows:

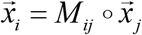

where *i* and *j* are the residue indices; *M*_ij_ is the rigid motion between two inter-domain residues *I* and *j*, 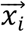 and 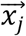 are the atomic coordinates in the local coordinate system of residue *i* and residue *j*, respectively.

### 3.1 Results of benchmark set

E2EDA was compared with SADA, DEMO and AIDA on the benchmark dataset. We assembled single-domain structures disassembled from native protein structures. For a fair comparison with other methods, homologous templates with a sequence identity >30% of the query were excluded.

The results of E2EDA, SADA, DEMO and AIDA on the benchmark set are summarised in Table 1. Meanwhile, detailed results for each protein are presented in Supplementary Table S4. On the benchmark set, the average TM-score of the model assembled by E2EDA is 0.84, which is 5.0% higher than that of SADA (0.80), 13.5% higher than that of DEMO (0.74) and 33.3% higher than that of AIDA (0.63) respectively. E2EDA correctly assembled 340 of the 356 targets, accounting for 95.5% of the total, 3.7%, 6.9% and 18.1% higher than SADA (328), DEMO (318) and AIDA (288), respectively. The *P*-values indicated that E2EDA was statistically significantly different from the other three methods. For different types of multi-domain proteins, Figure 2 visually compares the average TM-score of the four methods in box plots, indicating that the prediction network can effectively capture orientations between domains for domain assembly. Evidently, domain assembly became more difficult as the number of domains increased, as the degrees of freedom of domains increased with the number of domains. Moreover, correctly predicting multiple rigid motions is evidently more difficult than predicting one. For 2dis and 2dom, E2EDA had little improvement compared with SADA because the inter-domain orientation on 2dis and 2dom proteins was simpler and easier to predict making SADA already achieve relatively high accuracy and thus more difficult to improve. However, more importantly, on 3dom and m4dom, we improved significantly, which may have been due to E2EDA having fused some useful features through the attention mechanism to predict the inter-domain orientation accurately.

**Table 1.** Assembly results of E2EDA, SADA, DEMO and AIDA on benchmark proteins by using native domains. #TM-score ≥ 0.5 indicates the number of models with TM-score ≥ 0.5. *P* -value is the result of Wilcoxon signed-rank test based on comparison with the TM-score of E2EDA.

| Method | TM-score | | | | | #TM-score $\geq 0.5$ | $P$-value |
| --- | --- | --- | --- | --- | --- | --- | --- |
|  | 2dis | 2dom | 3dom | m4dom | Total |  |  |
| **E2EDA** | **0.91** | **0.86** | **0.79** | **0.68** | **0.84** | **340** | NA |
| SADA | 0.89 | 0.83 | 0.72 | 0.60 | 0.80 | 328 | 3.75E-06 |
| DEMO | 0.84 | 0.77 | 0.69 | 0.53 | 0.74 | 318 | 5.74E-25 |
| AIDA | 0.71 | 0.70 | 0.52 | 0.42 | 0.63 | 288 | 2.59E-80 |

**Figure. 2.**
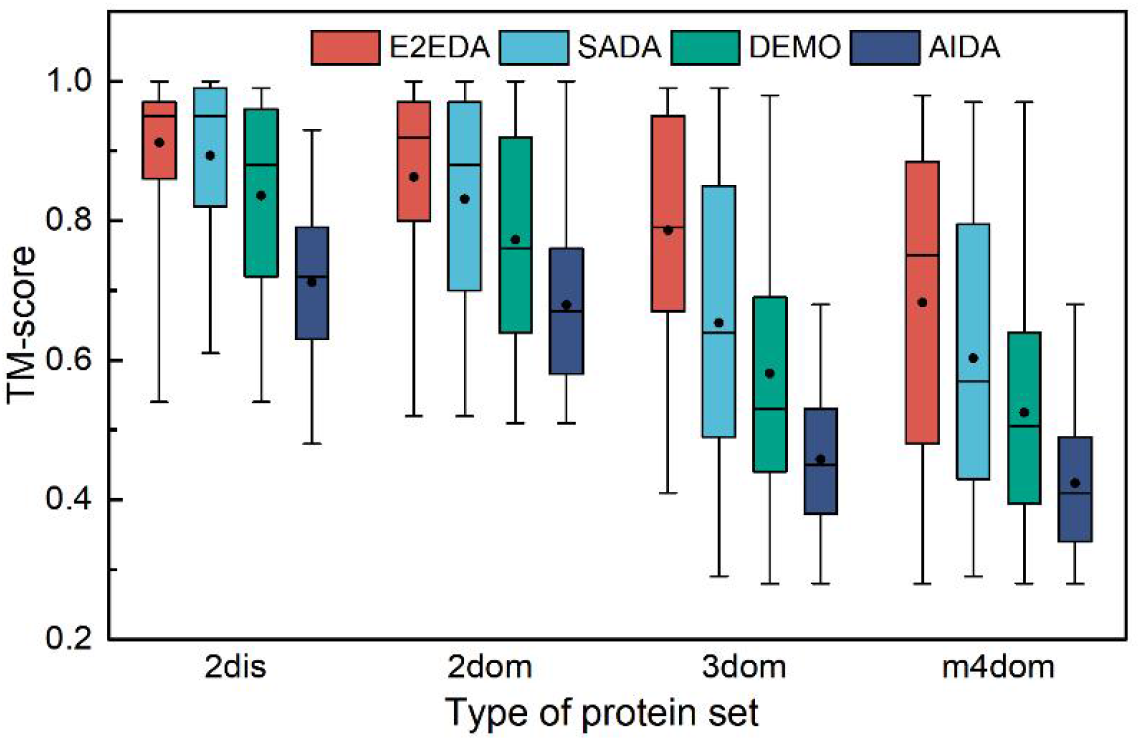
Boxplot for the TM-score of the assembly models by E2EDA, SADA, DEMO and AIDA. The square and horizontal lines in the box represent the mean and median TM-scores and the horizontal lines on the top and bottom are the maximum and minimum TM-scores, respectively.

### 3.2 Assembly human proteins predicted by AlphaFold2

Although many protein structures predicted by AlphaFold2 have reached experimental resolution, the training set of AlphaFold2 is heavily biased towards single-domain proteins, which makes the ability to capture inter-domain orientations relatively poor when performing full-chain modelling of multi-domain proteins [20]. E2EDA specialises in predicting the inter-domain orientation of multi-domain proteins for domain assembly. Therefore, the quality of the full-chain model predicted by AlphaFold2 may be further improved by E2EDA. To test the effect of the E2EDA reassembly of human proteins predicted by AlphaFold2, we constructed a human protein dataset. The human protein dataset had a total of 185 multi-domain proteins, all of which came from AlphaFold DB [28]. Moreover, the TM-score of the full-chain model was less than 0.8. These proteins had more than 70% coverage with native proteins and less than 70% sequence identity with each other. In this test, single-domain structures were disassembled directly from the full-chain protein structures in the AlphaFold DB.

A head-to-head comparison of the TM-score of the model reassembled by E2EDA and the full-chain model predicted by AlphaFold2 is shown in Figure 3A. Detailed results for each protein are presented in Supplementary Table S5. Among the 185 target proteins in the human protein dataset, the TM-score of 120 proteins increased after E2EDA reassembly, accounting for 64.9% of the total. The average TM-score increased from 0.59 to 0.63, an increase of 6.8%, which showed that E2EDA could effectively capture inter-domain orientation and improve the quality of the model by reassembling the full-chain model predicted by AlphaFold2.

**Figure. 3.**
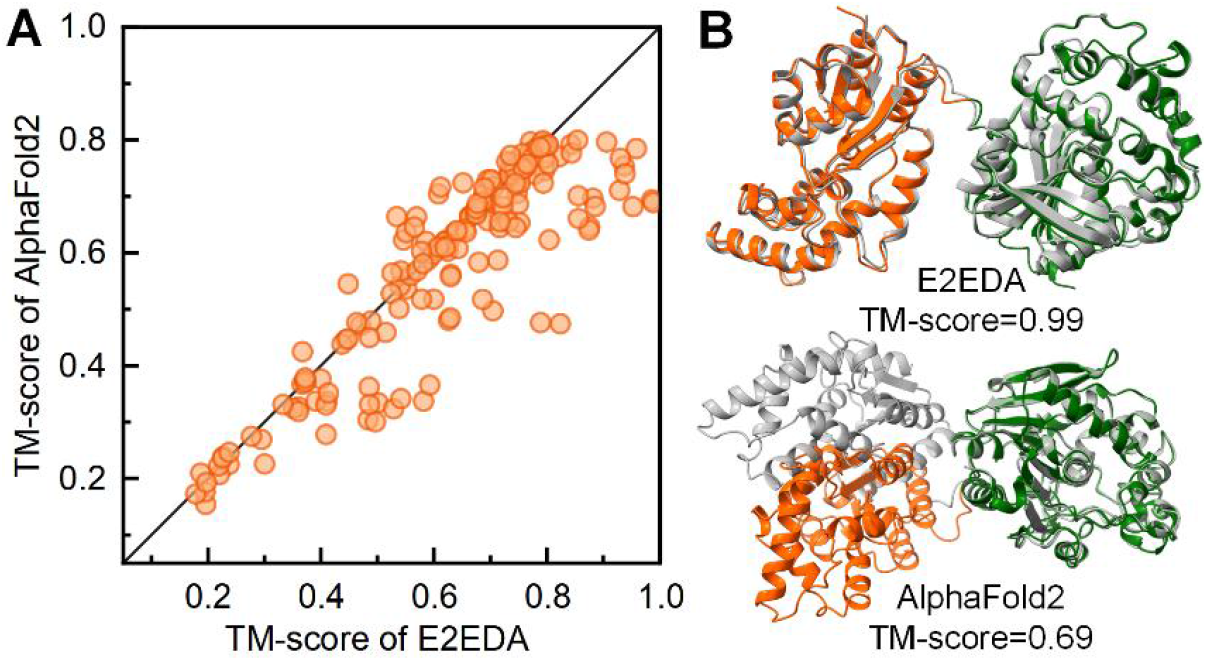
Comparison of E2EDA and AlphaFold2 in human protein dataset. (A) Head-to-head comparison between the full-chain model assembled by E2EDA and the full-chain model predicted by AlphaFold2. (B) Model of human protein 3wk4A predicted by AlphaFold2 and model after E2EDA reassembly. Orange and green indicate the two domains predicted by AlphaFold2, respectively, and grey indicates the native structure.

Figure 3B shows an example of improved model accuracy after E2EDA assembly. The protein 3wk4A consisted of two single-domain structures, and the TM-scores of domains 1 and 2 decomposed by the full-chain model predicted by AlphaFold2 were 0.99 and 1.0, respectively. The full-chain model predicted by AlphaFold2 was only 0.69, which indicated that AlphaFold2 might have been able to predict the single-domain structure accurately. However, it had more difficulty capturing the orientation information between domains. The TM-score of the full-chain model reassembled by E2EDA was 0.99, which was 43.5% higher than the full-chain model predicted by AlphaFold2 (0.69), indicating that E2EDA could capture the correct inter-domain orientation to assemble a higher-quality model.

### 3.3 Results of CASP14 targets

E2EDA was tested on 17 multi-domain proteins from the CASP14 official website (https://predictioncenter.org/download_area/CASP14/) and compared with the state-of-the-art method AlphaFold2. In this section, we conducted two sets of experiments using different single-domain structures. In the first set of experiments, single-domain structures were disassembled from the full-chain structure predicted by AlphaFold2 of CASP14. In the second set of experiments, the single-domain structure was decomposed from the experimental structure.

Detailed results for each target are in Supplementary Table S6 and intuitively presented in Figure 4A. In the first set of experiments, E2EDA reassembled the model predicted by AlphaFold2. The average TM-score of the model reassembled by E2EDA was 0.829, which was slightly higher than the full-chain model predicted by AlphaFold2 (0.822). In the second set of experiments, E2EDA assembled the single-domain model decomposed from the experimental structure into a full-chain model, with an average TM-score of 0.872, which was 6.1% higher than the full-chain model predicted by AlphaFold2 (0.822). The comparison of CSAP14 results showed that E2EDA could improve the accuracy of the model by reassembling the model predicted by AlphaFold2. If a higher-quality single-domain model could be obtained, then the quality of the full-chain model would be further improved. Figure 4B and D shows a target protein T1024, which improved the quality of the full-chain model predicted by AlphaFold2 after assembly by E2EDA. For the target T1024, the full-chain model predicted by AlphaFold2 consisted of two single-domain domains with TM-scores of 0.96 and 0.98, respectively, whereas the TM-score of the full-chain model was only 0.71. In the Predicted Aligned Error (PAE) plots of Figure 4C and E [28], the colour of the inter-domain filling of the model predicted by AlphaFold2 is evidently lighter than that of the model assembled by E2EDA, which indicated that the inter-domain orientation of the model assembled by E2EDA was more accurate than that of AlphaFold2.

**Figure. 4.**
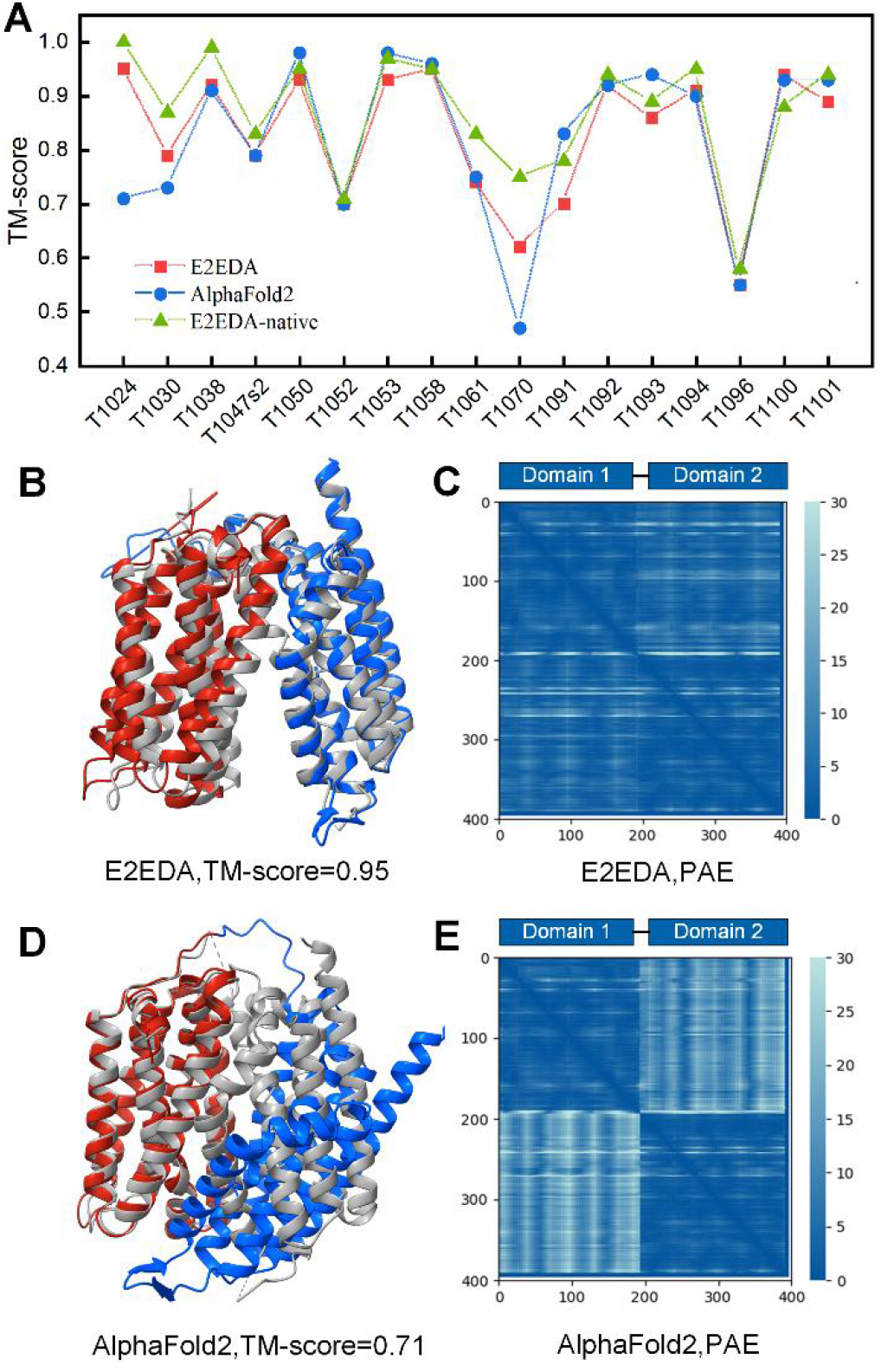
Comparing the performance of E2EDA and AlphaFold2 on 17 multi-domain proteins of CASP14. (A) TM-score of models predicted by AlphaFold2, E2EDA and E2EDA-native. (B) and (D) are the full-chain models of protein T1024 predicted by E2EDA and AlphaFold2, respectively. Different colour represent different domains, and gray represents the native structure. (C) and (E) are the predicted aligned error (PAE) plots of the model predicted by E2EDA and AlphaFold2 and the native structure, respectively.

### 3.4 Analysis of end-to-end assembly efficiency

To verify whether the end-to-end assembly method can improve the assembly efficiency, we compared the running time of E2EDA and SADA on the benchmark set under the given hardware (CPU: Intel(R) Xeon(R) Gold 6226R CPU @ 2.90GHz, GPU: V100 with 32GB), which was convenient for comparison because SADA was also developed by us. The running times of E2EDA and SADA on the benchmarks are summarised in Supplementary Table S7, and detailed results for each protein can be found in Supplementary Table S8. For an intuitive comparison of the running time between E2EDA and SADA, their results are depicted in Figure 5. The average running time of E2EDA was reduced by 74.6% compared with SADA. E2EDA obtained shorter running time on 328 out of 356 proteins, accounting for 92.1% of the total proteins. The results showed that the end-to-end assembly method could effectively improve the assembly efficiency compared with the simulation-based domain assembly method because the end-to-end assembly method could directly predict the orientation of the domains for assembly without a time-consuming simulation process. Furthermore, recently, an alternative to MSA-based models has emerged –leveraging information contained in pre-trained protein language models (pLMs) [29-31]. These pLMs are trained on huge sequence databases, and the internal representations (embeddings) learned are successfully applied to many downstream tasks [32-34]. Therefore, replacing MSA with pLM embedding can reduce the time of searching for MSA [35] and further improve the assembly efficiency of E2EDA.

**Figure. 5.**
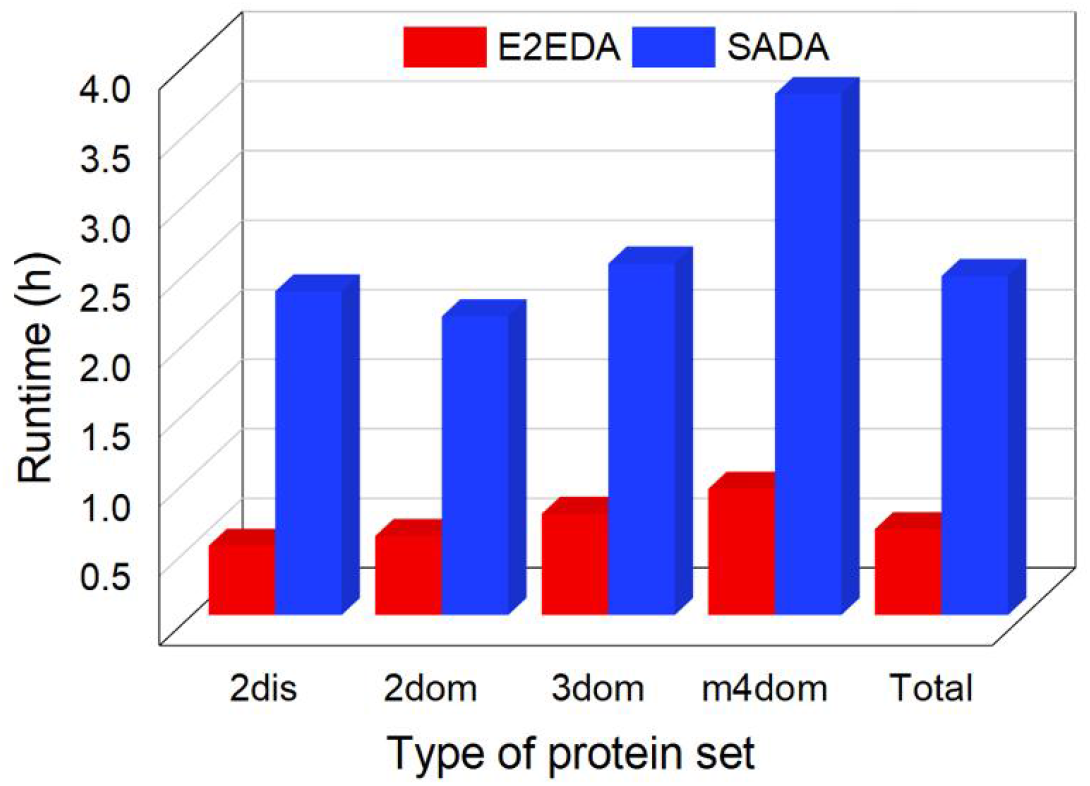
Comparison of the average running time of E2EDA and SADA on different types of proteins in the benchmark set.

Although the structures of many protein targets predicted by AlphaFold2 are at or close to the experimental resolution, it has a large demand for hardware resources. Moreover, it easily exceeds the memory of a single GPU when inferring large proteins. For a V100 with 16GB of memory, AlphaFold2 can predict proteins with a maximum of about 1,300 residues. GPU memory usage is roughly quadratic in the number of residues [18]. Hence, the memory required to predict a 2,000-residue protein can greatly exceed that of a single V100. In E2EDA, for large proteins, based on the divide-and-conquer strategy, we first predict the single-domain structure and then assemble them into a full-chain model, which can effectively reduce the demand for hardware resources.

### 3.5 Ablation study

To analyse the impact of MSA features, template features and single-domain features on the performance of E2EDA, we constructed two versions different from E2EDA: E2EDA-w/o-S and E2EDA-w/o-S&T. We trained E2EDA-w/o-S and E2EDA-w/o-S&T with the same parameters and training set as E2EDA, where E2EDA-w/o-S had no single-domain features and E2EDA-w/o-S&T had no single-domain features and templates feature.

The results of E2EDA, E2EDA-w/o-S and E2EDA-w/o-S&T on the benchmark set are summarised in Table 2. Detailed results for each protein can be found in Supplementary Table S9. As shown in Table 2, E2EDA outperforms E2EDA-w/o-S and E2EDA-w/o-S&T on all types of protein sets in the benchmark set. The average TM-score of E2EDA on the benchmark protein was 0.84, 10.5% and 2.4% higher than E2EDA-w/o-S&T (0.76) and E2EDA-w/o-S (0.82), respectively. In the significance test results, the *P*-values of E2EDA-w/o-S and E2EDA-w/o-S&T were 8.9E-11 and 8.4E-18, respectively. These outcomes indicated that the performance of E2EDA was significantly better than that of E2EDA-w/o-S and E2EDA-w/o-S&T, showing the effectiveness of template features and single-domain features.

**Table 2.** The assembly results of E2EDA, E2EDA-w/o-S and E2EDA-w/o-S&T on the benchmark set, where E2EDA-w/o-S had no single-domain features and E2EDA-w/o-S&T had no single-domain features and template features.

| Method | TM-score | | | | | #TM-score $\geq 0.8$ | *P*-value |
| --- | --- | --- | --- | --- | --- | --- | --- |
|  | 2dis | 2dom | 3dom | m4dom | Total |  |  |
| **E2EDA** | **0.91** | **0.86** | **0.79** | **0.68** | **0.84** | **246** | NA |
| E2EDA-w/o-S | 0.89 | 0.84 | 0.76 | 0.66 | 0.82 | 220 | 8.9E-11 |
| E2EDA-w/o-S&T | 0.84 | 0.79 | 0.69 | 0.59 | 0.76 | 173 | 8.4E-18 |

Figure 6 compares the quality of the models assembled by the three methods more intuitively, thereby showing that both single-domain features and template features could effectively improve the accuracy of the assembled models. Compared with single-domain features, template features were evidently more effective. For protein 2ahvA in Figure 6E, the TM-score of the predicted structures of E2EDA-w/o-S&T without template information and E2EDA-w/o-S with template information were 0.55 and 0.96, respectively. Figure 6F are the two templates used during protein 2ahvA assembly. Although the TM-score of the two templates was only 0.77 and 0.75, respectively, they both contained the correct inter-domain orientation, which made the TM-score of the assembled model reach 0.96, indicating that E2EDA could capture the correct inter-domain orientation from multiple templates to improve assembly accuracy. The accuracy of some proteins also decreased after adding template information. For the 2d1cA protein in Supplementary Figure S4A, the TM-score of the predicted structure dropped from 1.0 to 0.9 after adding template information. As shown in Supplementary Figure S4B, the template did not contain inter-domain position information, thus affecting the prediction accuracy.

**Figure. 6.**
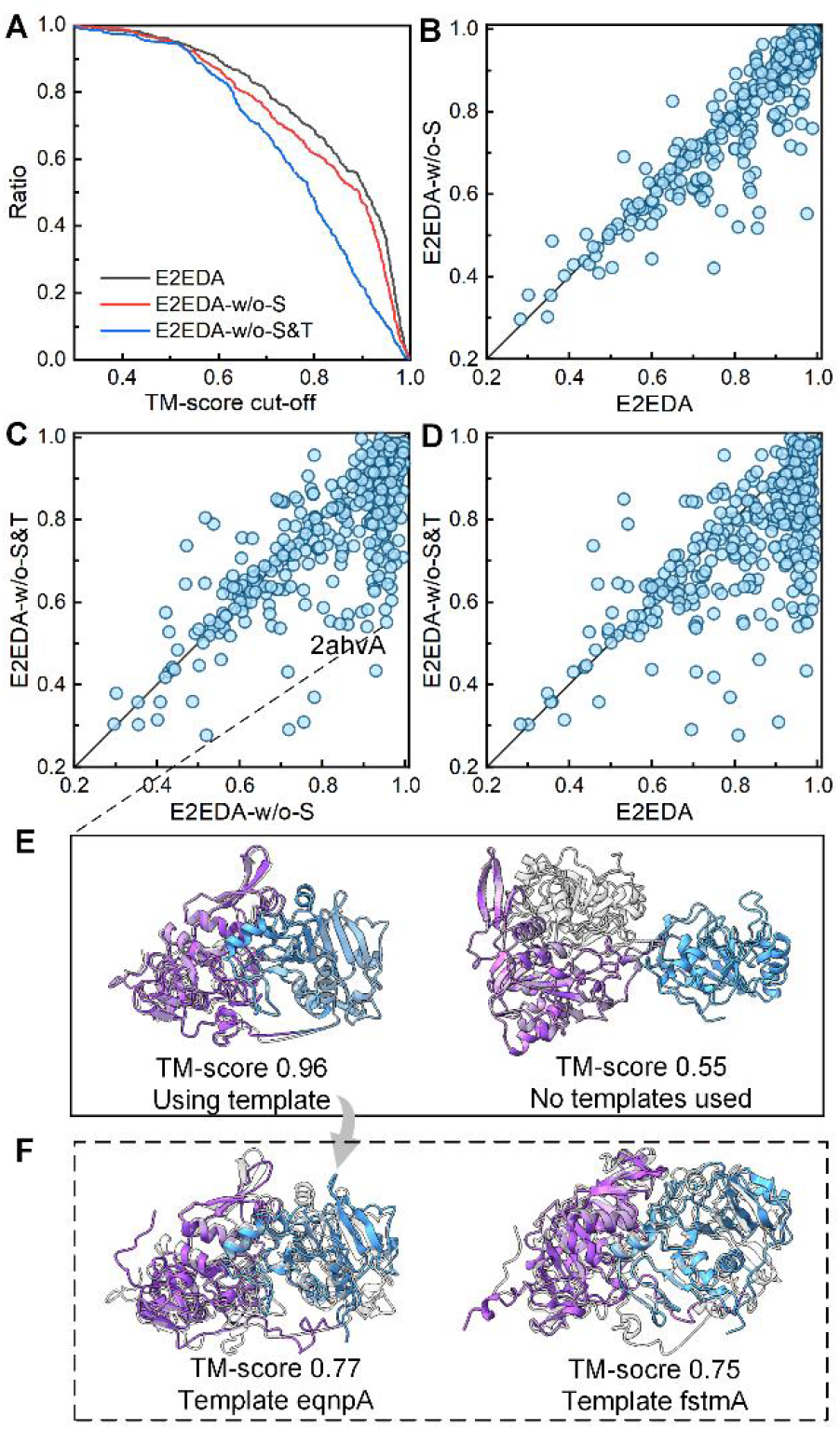
Comparison between E2EDA, E2EDA-w/o-S and E2EDA-w/o-S&T, where E2EDA-w/o-S had no single-domain features, and E2EDA-w/o-S&T had no single-domain features and templates feature. (A) is a line chart of the ratio of the number of models above the TM-score cut-off threshold to the total number. (B), (C) and (D) are TM-score head-to-head comparisons among the three methods. (E) are full-chain models assembled from template-based E2EDA-w/o-S and non-template-based E2EDA-w/o-S&T, respectively. Purple and blue represent different domains, respectively, and grey represents native structures. (F) Top two templates of protein 2ahvA searched by HHsearch.

### 3.6 Effect of using RMscore

In E2EDA, the 50 rigid motions with the highest confidence are used to assemble 50 models, which are scored by RMscore to select the top five models as the final model or the single-domain structure for the next assembly. To verify the effect of RMscore, the complete E2EDA method is compared with E2EDA without RMscore (E2EDA-w/o-R) on the benchmark set. Here, E2EDA-w/o-R indicates that RMsocre is not used to score the assembled structures and that the final model is the one assembled using the rigid motion with the highest confidence.

Supplementary Table S10 summarises the assembly results of E2EDA and E2EDA-w/o-R on the benchmark. Detailed results for each protein can be found in Supplementary Table S11. Compared with E2EDA-w/o-R (0.72), the average TM-score of E2EDA (0.84) increased by 16.7%. The average TM-score of E2EDA was 15.2% and 11.7% higher than that of E2EDA-w/o-R on 2dom protein and 3dom protein, respectively. Especially on 3dom protein and m4dom protein, the average TM-score of E2EDA was 21.5% and 25.9% higher than that of E2EDA-w/o-R, respectively. Figure 7A shows the histogram of the average TM-score on different types of multi-domain proteins, which compares the effect of RMscore more intuitively.

**Figure. 7.**
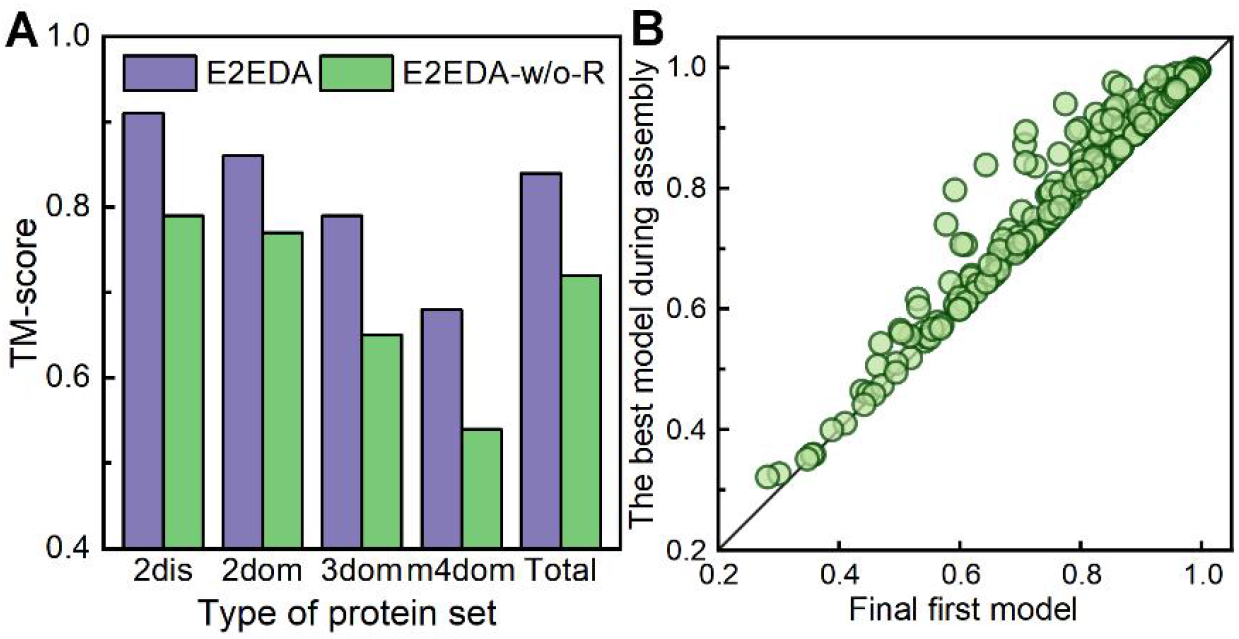
Test the performance of RMscore on benchmark proteins. (A) Average TM-score of models assembled by E2EDA and E2EDA-w/o-R, where E2EDA-w/o-R indicates that E2EDA does not use the scoring strategy RMscore (B) TM-score head-to-head comparison between the best model during E2EDA assembly and the final first model selected by RMscore.

To further verify the effect of RMscore further, we compared the TM-score between the best model during E2EDA assembly process and the final first model on the benchmark set. These results are summarised in Supplementary Table S12 and detailed results for each protein can be found in Supplementary Table S13. The average TM-score of the final first model (0.84) was reduced by 1.2% compared with the best model during assembly (0.85). A head-to-head comparison of the TM-score of the best model during assembly and the TM-score of the final first model is shown in Figure 7B. E2EDA assembled 294 out of 356 targets when the error of the two TM-score was less than 0.03, accounting for 82.6% of the total. The above experimental results showed that RMscore could effectively improve the quality of the model assembled by E2EDA.

## 4 Conclusion

We developed a deep learning-based end-to-end domain assembly method, E2EDA, which improves the quality of assembled models while improving assembly efficiency. In E2EDA, a neural network is first constructed to predict the rigid motion between residues of multi-domain proteins. In addition to the MSA information, the attention mechanism is also used to fuse the information of the single-domain domain and the information of multiple templates to improve the accuracy of rigid motion prediction between residues. As the confidence and accuracy of the predicted rigid motions do not completely correspond, the 50 rigid motions with the highest confidence will be used to assemble 50 models. The best model will be selected through the scoring strategy RMscore, thus improving the assembly accuracy.

E2EDA was compared with SADA, DEMO and AIDA on 356 benchmark proteins. The experimental results showed that E2EDA outperformed the other three protein domain assembly methods. Meanwhile, compared with SADA, the average running time of E2EDA on the benchmark set was reduced by 74.6%, which was due to the ability of E2EDA to predict inter-domain orientations and perform domain assembly directly without the assembly simulation process. Experimental results on human protein datasets and multi-domain proteins from CASP14 show that E2EDA can capture more accurate inter-domain orientations than AlphaFold2 and can reassemble higher-quality full-chain models. In addition, replacing MSA with pre-trained protein language model embeddings [29-31, 35] to improve assembly efficiency and attempting to extend the E2EDA-based method to the assembly of complexes are also the direction of our follow-up exploration.

## 5 Acknowledgments

This work is supported by the “New Generation Artificial Intelligence” major project of Science and Technology Innovation 2030 of the Ministry of Science and Technology of the People’s Republic of China [No. 2022ZD0115100], the National Nature Science Foundation of China [No. 62173304], and the Key Project of Zhejiang Provincial Natural Science Foundation of China [No. LZ20F030002].

