## Supplementary information for "E2EDA: Protein domain assembly based on end-to-end deep learning"

#### Supplementary Methods

**Method S1.** The construction process of the rigid motion $M_i$ from the local coordinate system to the global coordinate system of the *i*-th residue. We use N, $C_\alpha$, and C three atoms to construct the local coordinate system of the *i*-th residue.

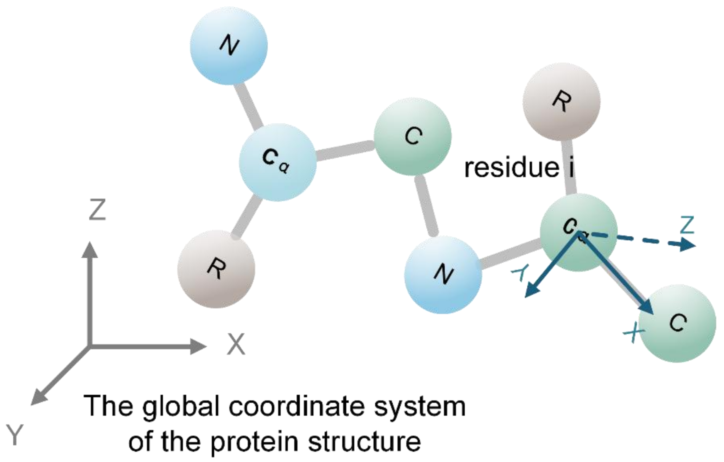


def Method1 $(\vec{x}_N, \vec{x}_{C_\alpha}, \vec{x}_C)$ :

1: $\vec{v}_1 = \vec{x}_C - \vec{x}_{C_\alpha}$

2: $\vec{v}_2 = \vec{x}_N - \vec{x}_{C_\alpha}$

3: $\vec{e}_1 = \vec{v}_1 / \|\vec{v}_1\|$

4: $\vec{u}_2 = \vec{v}_2 - \vec{e}_1 \left(\vec{e}_1^{\mathrm{T}} \vec{v}_2\right)$

5: $\vec{e}_2 = \vec{u}_2 / \|\vec{u}_2\|$

6: $\vec{e}_3 = \vec{e}_1 \times \vec{e}_2$

7: $R = \mathrm{concat}\left(\vec{e}_1, \vec{e}_2, \vec{e}_3\right)$

8: $Q = \mathrm{rotateMatrixToQuaternion}\,(R)$

9: $M_i = \vec{x}_{C_\alpha}$

return $M_i$

def rotateMatrixToQuaternion$(R)$:

$w = \sqrt{r_{11} + r_{22} + r_{33} + r_{44} + 1} / 2$

$x = (r_{23} - r_{32}) / 4w$

$y = (r_{31} - r_{13}) / 4w$

$z = (r_{12} - r_{21}) / 4w$

$Q = (w, x, y, z)$

return $Q$

Where $\vec{e}_1$, $\vec{e}_2$ and $\vec{e}_3$ represent the three basis vectors of the constructed local coordinate system; $R$ represents the rotation matrix transformed from the local coordinate system to the global coordinate system; $r_{ij}$ is the value of the corresponding position of the rotation matrix $R$. $w$, $x$, $y$ and $z$ are the four components of the quaternion $Q$.

**Method S2.** Detailed calculation steps for the rigid motion $M_{ij}$ between the $i$-th residue and the $j$-th residue. We denote the application of rigid motions to atomic positions by the ◦ operator:

$$\begin{aligned} \vec{x}_{global} &= M_i \circ \vec{x}_i \\ &= (Q_i, T_i) \circ \vec{x}_i \\ &= (R_i, T_i) \circ \vec{x}_i \\ &= R_i \vec{x}_i + T_i \end{aligned}$$

We also use the ◦ operator to denote combinations of rigid motions:

$$\begin{aligned} M_{\text{result}} &= M_i \circ M_j \\ &= (Q_i, T_i) \circ (Q_j, T_j) \\ &= (R_i, T_i) \circ (R_j, T_j) \\ &= (R_i R_j, R_i T_j + T_i) \end{aligned}$$

In addition, the inverse of the rigid motion $M_i$ is $M_i^{-1}$:

$$\begin{aligned} M_i^{-1} &= (Q_i, T_i)^{-1} \\ &= (Q_i^{-1}, -Q_i^{-1} T_i) \\ &= (R_i^{-1}, -R_i^{-1} T_i) \end{aligned}$$

The calculation process of the rigid motion $M_{ij}$ between the $i$-th residual and the $j$-th residual is as follows:

$$\begin{aligned} M_{ij} &= M_i^{-1} \circ M_j \\ &= (Q_i^{-1}, -Q_i^{-1} T_i) \circ (Q_j, T_j) \\ &= (Q_i^{-1} Q_j, Q_i^{-1} T_j - Q_i^{-1} T) \\ &= (R_i^{-1} R_j, R_i^{-1} T_j - R_i^{-1} T) \end{aligned}$$

**Method S3.** The quaternion of rigid motion is used to construct the rotation matrix between two domains, which can be calculated as follows:

$$R_i = \begin{pmatrix} w_i^2 - x_i^2 - y_i^2 + z_i^2 & 2(x_i y_i - w_i z_i) & 2(x_i z_i + w_i y_i) \\ 2(x_i y_i + w_i z_i) & w_i^2 - x_i^2 + y_i^2 - z_i^2 & 2(y_i z_i - w_i x_i) \\ 2(x_i z_i - w_i y_i) & 2(y_i z_i + w_i x_i) & w_i^2 - x_i^2 - y_i^2 + z_i^2 \end{pmatrix}$$

where $w_i$, $x_i$, $y_i$, and $z_i$ are the four scalars of the unit quaternion.

### Supplementary Figures

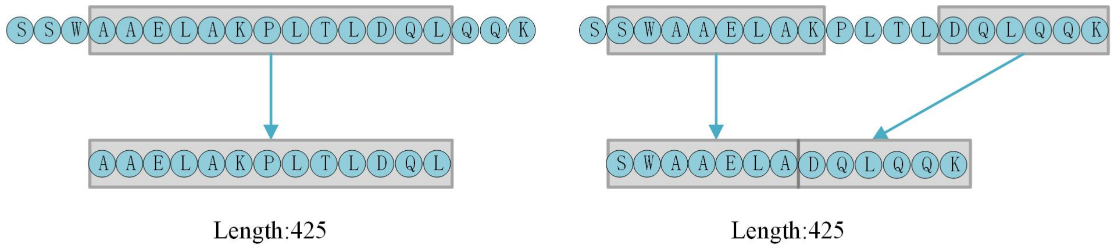


**Figure S1.** To reduce the loss of large protein information, two subsequences were randomly selected, one from the first half of the original sequence and the other from the second half, and then spliced together to form a new sequence.

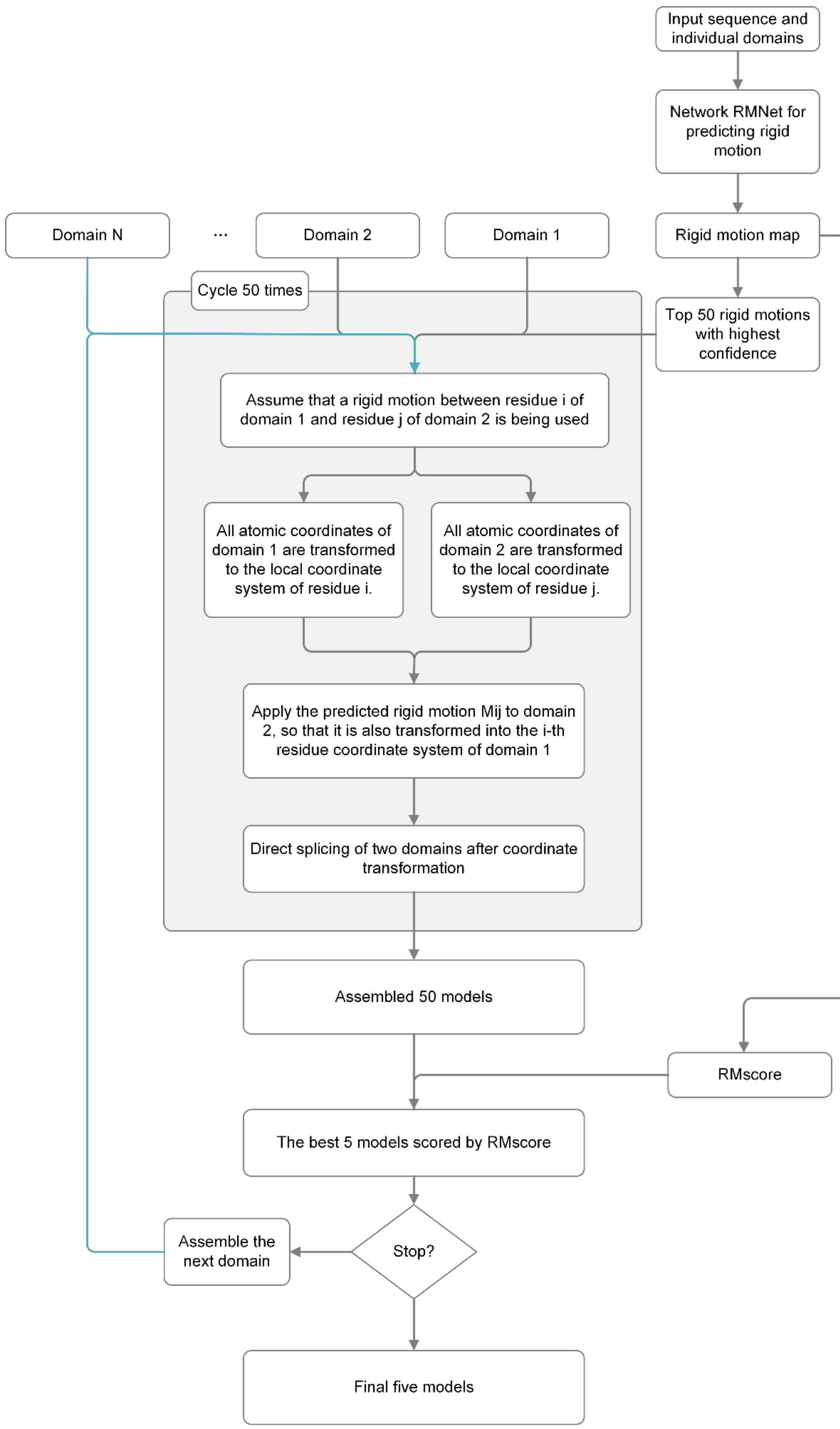


**Figure S2.** The detailed process of domain assembly in E2EDA, the blue line in the figure indicates that it is not input at the same time as the gray line.

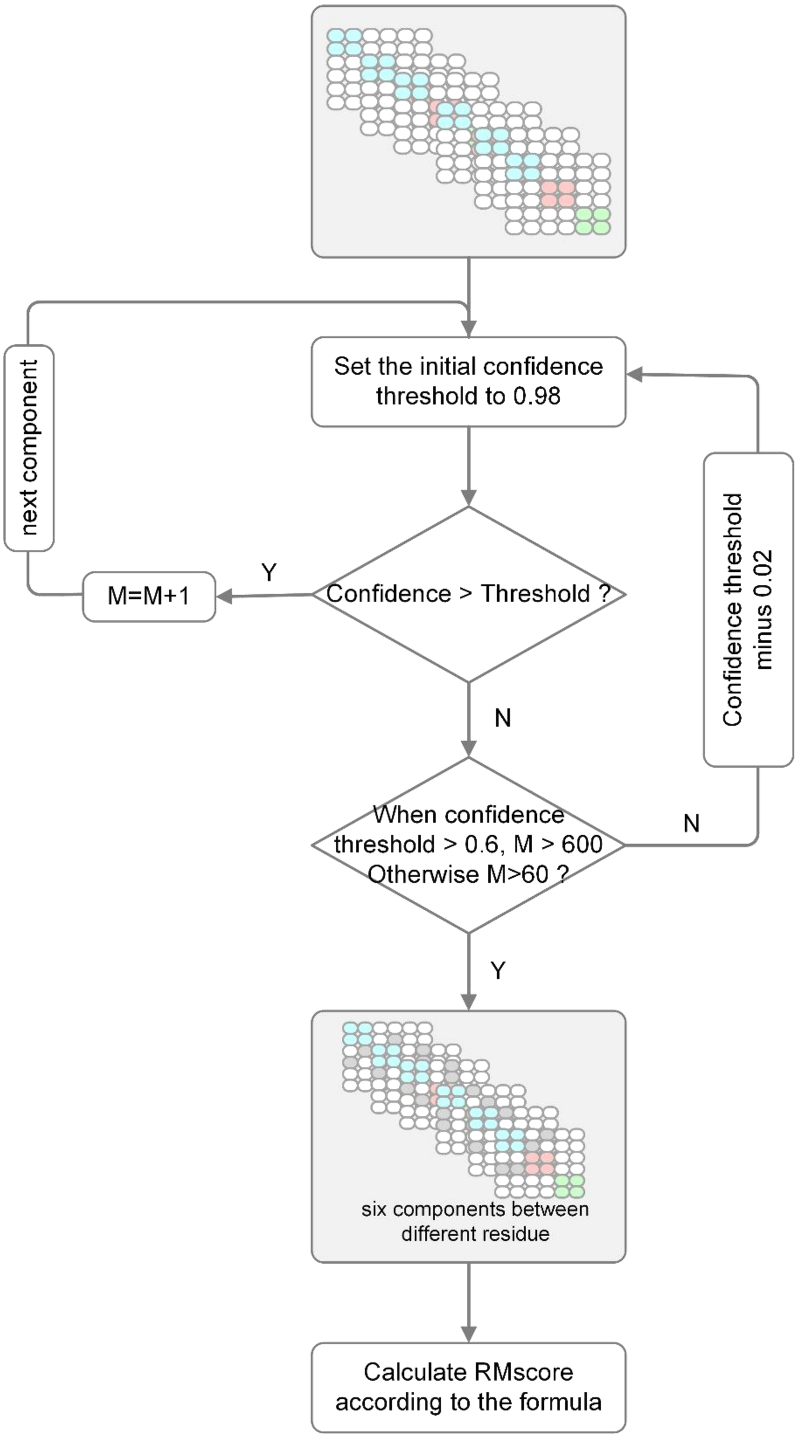


**Figure S3.** The detailed process of constructing the RMscore.

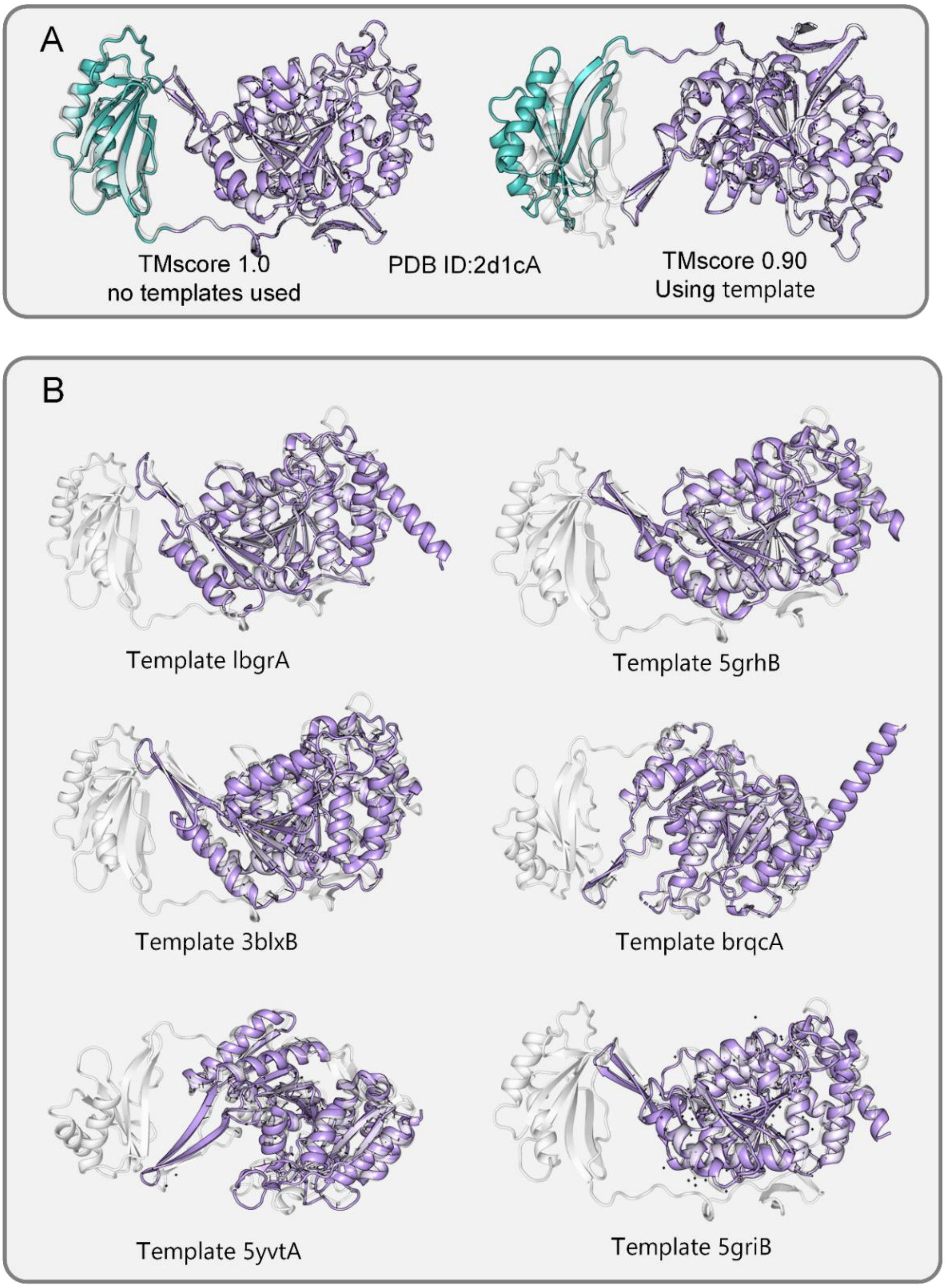


**Figure S4.** Blue and purple represent two different domains, respectively, and gray represents the native structure (A) For protein 2d1cA, models assembled with templates and models assembled without templates. (B) The searched template of protein 2d1cA, which has no blue domain, indicating that there is only one purple domain structure in the template.

### Supplementary Tables

**Table S1.** All features used by E2EDA

| type | name | shape |
|---|---|---|
| sequence features | one-hot encoding of the input sequence | $L$×20 |
| | location specifficity frequency matrix | $L$×21 |
| | positional entropy | $L \times 1$ |
| | mean product correction | $L \times L \times 1$ |
| | coupling matrix | $L \times L \times 442$ |
| single-domain features | domain structure information | $L \times L \times 6$ |
| | domain boundary information | $L \times L \times 9$ |
| template features | multiple template structure information | $N \times L \times L \times 6$ |

**Table S2.** Detailed parameters of Fused-MBConv and MBConv blocks.

| Stage | Operator | Kernel | Stride | Expansion | In_c | Out_c | Layers |
|---|---|---|---|---|---|---|---|
| 1 | Fused-MBConv | 3 | 1 | 1 | 64 | 64 | 2 |
| 2 | Fused-MBConv | 3 | 1 | 4 | 64 | 64 | 4 |
| 3 | Fused-MBConv | 3 | 1 | 4 | 64 | 128 | 4 |
| 4 | MBConv | 3 | 1 | 4 | 128 | 128 | 6 |
| 5 | MBConv | 3 | 1 | 6 | 128 | 128 | 9 |
| 6 | MBConv | 3 | 1 | 6 | 128 | 128 | 15 |

**Table S3.** Information for each protein in the benchmark dataset. #Domains indicates the number of domains of the protein. 2dis indicates that the protein has 2 discontinuous domains.

| PDB ID | #Domains | PDB ID | #Domains | PDB ID | #Domains | PDB ID | #Domains |
|---|---|---|---|---|---|---|---|
| 1cjyA | 2 | 3isqA | 2 | 2b5uA | 3 | 4fe9A | 4 |
| 1efdN | 2 | 3isqA | 2 | 2ewfA | 3 | 4h2aA | 4 |
| 1fjrA | 2 | 3k1rA | 2 | 2piaA | 3 | 4i5sB | 4 |
| 1g87B | 2 | 3k2iA | 2 | 2r7dA | 3 | 4iggB | 6 |
| 1hx6B | 2 | 3kh5A | 2 | 2uwnA | 3 | 4j9vA | 4 |
| 1iwaA | 2 | 3kjpA | 2 | 2v0nA | 3 | 4k3bA | 6 |
| 1m5qH | 2 | 3kt1A | 2 | 2vgmA | 3 | 4kwuA | 6 |
| 1mkfA | 2 | 3ktmE | 2 | 2wqrB | 3 | 4m00A | 4 |
| 1mkmB | 2 | 3kzwA | 2 | 2y25B | 3 | 1bp1A | 2dis |
| 1nh2D | 2 | 3l76A | 2 | 2yk0A | 3 | 1cjsA | 2dis |
| 1pprM | 2 | 3ld1A | 2 | 2zzqA | 3 | 1ck1A | 2dis |
| 1prrA | 2 | 3lsgA | 2 | 3bt1U | 3 | 1ecrA | 2dis |
| 1q19A | 2 | 3me4A | 2 | 3c1yA | 3 | 1f5qD | 2dis |
| 1qwrA | 2 | 3ml4C | 2 | 3cw2C | 3 | 1fa9A | 2dis |
| 1r71B | 2 | 3mx2B | 2 | 3f83A | 3 | 1gu7A | 2dis |
| 1rh1A | 2 | 3mzfA | 2 | 3fc3A | 3 | 1itwA | 2dis |
| 1rktA | 2 | 3njaB | 2 | 3gbgA | 3 | 1jkiA | 2dis |
| 1s6lA | 2 | 3nqiA | 2 | 3h5cB | 3 | 1n80A | 2dis |
| 1sp3A | 2 | 3nt8A | 2 | 3ibjA | 3 | 1nzjA | 2dis |
| 1vz6A | 2 | 3og5A | 2 | 3ippB | 3 | 1qhdA | 2dis |
| 1w3aA | 2 | 3oh0A | 2 | 3jymB | 3 | 1qz9A | 2dis |
| 1wv3A | 2 | 3pcsB | 2 | 3kbgA | 3 | 1sb7B | 2dis |
| 1x7pA | 2 | 3po3S | 2 | 3mc8A | 3 | 1vk1A | 2dis |
| 1x9yA | 2 | 3pxpA | 2 | 3npfA | 3 | 1vrmA | 2dis |
| 1y11A | 2 | 3qavA | 2 | 3orjA | 3 | 1xvuA | 2dis |
| 1yiqA | 2 | 3qf4B | 2 | 3plaA | 3 | 1yy3A | 2dis |
| 1zbuB | 2 | 3qjjA | 2 | 3qe9Y | 3 | 1z87A | 2dis |
| 1ze1A | 2 | 3qtdA | 2 | 3qjoA | 3 | 2a1sC | 2dis |
| 2ablA | 2 | 3r6bA | 2 | 3qphA | 3 | 2a3lA | 2dis |
| 2ahvA | 2 | 3rh7A | 2 | 3qyeA | 3 | 2bt1A | 2dis |
| 2bkpA | 2 | 3rwxA | 2 | 3rimA | 3 | 2bydA | 2dis |
| 2c1yA | 2 | 3sb4A | 2 | 3rrpA | 3 | 2c43A | 2dis |
| 2cxcA | 2 | 3swjA | 2 | 3soaA | 3 | 2dfyC | 2dis |
| 2d1cA | 2 | 3t58B | 2 | 3tixD | 3 | 2dlaA | 2dis |
| 2d7iA | 2 | 3t7jA | 2 | 3tp9A | 3 | 2g3pA | 2dis |
| 2e9hA | 2 | 3u07C | 2 | 3ua3A | 3 | 2gg6A | 2dis |
| 2e9xB | 2 | 3u0oB | 2 | 3uj0A | 3 | 2gsyE | 2dis |
| 2evrA | 2 | 3u9gA | 2 | 3vn4A | 3 | 2gzoA | 2dis |
| 2ew9A | 2 | 3ub1D | 2 | 3vsmA | 3 | 2j2cA | 2dis |
| 2fd5A | 2 | 3uitD | 2 | 3w1bA | 3 | 2kfwA | 2dis |
| 2gh8A | 2 | 3uo3A | 2 | 3zh9B | 3 | 2l9yA | 2dis |
| 2gt1A | 2 | 3v7oB | 2 | 4alzA | 3 | 2ntyB | 2dis |
| 2gzaC | 2 | 3vr8B | 2 | 4ax8A | 3 | 2r3vA | 2dis |
| 2hjqA | 2 | 3wkuA | 2 | 4b3iA | 3 | 2r58A | 2dis |
| 2hwjA | 2 | 3zvmA | 2 | 4bd9B | 3 | 2w4mA | 2dis |
| 2ijd1 | 2 | 4acoA | 2 | 4c0aB | 3 | 2x0cA | 2dis |
| 2iu7A | 2 | 4ap5A | 2 | 4c0sA | 3 | 2y51A | 2dis |
| 2iw2A | 2 | 4axdA | 2 | 4dimA | 3 | 2yb0E | 2dis |

| PDB ID | #Domains | PDB ID | #Domains | PDB ID | #Domains | PDB ID | #Domains |
|---|---|---|---|---|---|---|---|
| 2jz4A | 2 | 4bfiB | 2 | 4indA | 3 | 2z86C | 2dis |
| 2kdyA | 2 | 4bt9B | 2 | 4jdzB | 3 | 3afoA | 2dis |
| 2kn4A | 2 | 4cczA | 2 | 4kc3B | 3 | 3bu2A | 2dis |
| 2mbgA | 2 | 4d0nB | 2 | 4kikB | 3 | 3cvzA | 2dis |
| 2nsfA | 2 | 4d1iG | 2 | 4lmfA | 3 | 3dupA | 2dis |
| 2nykA | 2 | 4dj3A | 2 | 4lziA | 3 | 3eswA | 2dis |
| 2o6yA | 2 | 4dqaA | 2 | 4m9pA | 3 | 3eukH | 2dis |
| 2owbA | 2 | 4eo3A | 2 | 4pt5A | 3 | 3fi7A | 2dis |
| 2qfiA | 2 | 4eogA | 2 | 4uwhA | 3 | 3fvvA | 2dis |
| 2qp2A | 2 | 4etxA | 2 | 1c1zA | 5 | 3gmsA | 2dis |
| 2qygA | 2 | 4fguA | 2 | 1d2pA | 4 | 3hzzB | 2dis |
| 2r5wB | 2 | 4fkcA | 2 | 1k7tA | 4 | 3m1uA | 2dis |
| 2uu7A | 2 | 4fxkC | 2 | 1kfqA | 4 | 3mw8A | 2dis |
| 2w4bA | 2 | 4gbyA | 2 | 1ldjA | 7 | 3mwcA | 2dis |
| 2x7iA | 2 | 4ggmX | 2 | 1nyqB | 4 | 3nsjA | 2dis |
| 2x8kC | 2 | 4gslA | 2 | 1ug9A | 4 | 3ntkA | 2dis |
| 2yilA | 2 | 4gyjA | 2 | 1z1wA | 4 | 3oaaG | 2dis |
| 2yrqA | 2 | 4h3tA | 2 | 2au3A | 4 | 3ptyA | 2dis |
| 2zxcA | 2 | 4hmoA | 2 | 2ii2A | 4 | 3rfyA | 2dis |
| 3a1iA | 2 | 4ie6A | 2 | 2olsA | 4 | 3seoB | 2dis |
| 3a45A | 2 | 4l5gA | 2 | 2ra1A | 5 | 3spgA | 2dis |
| 3a56A | 2 | 4lpqA | 2 | 2v5dA | 4 | 3u0kA | 2dis |
| 3ajvA | 2 | 4m8rA | 2 | 2xt6A | 4 | 3vlaA | 2dis |
| 3aqkA | 2 | 4n06B | 2 | 2zpaB | 4 | 3vstA | 2dis |
| 3arbA | 2 | 4nj5A | 2 | 3apoA | 7 | 4aqfB | 2dis |
| 3aujG | 2 | 4opaB | 2 | 3b43A | 6 | 4b21A | 2dis |
| 3b2zF | 2 | 4qkuB | 2 | 3gf5B | 7 | 4dt4A | 2dis |
| 3b7wA | 2 | 4up9A | 2 | 3hjlA | 4 | 4dtfA | 2dis |
| 3bt3A | 2 | 4w7sA | 2 | 3kq4B | 4 | 4ewtA | 2dis |
| 3c4tA | 2 | 1bf2A | 3 | 3kwlA | 4 | 4f23A | 2dis |
| 3craA | 2 | 1bhgA | 3 | 3ob8A | 5 | 4fzbC | 2dis |
| 3d30A | 2 | 1f7uA | 3 | 3opfB | 4 | 4g1pA | 2dis |
| 3eo5A | 2 | 1fx7A | 3 | 3p53A | 4 | 4gfqA | 2dis |
| 3errA | 2 | 1griA | 3 | 3pvlA | 5 | 4hvzA | 2dis |
| 3g79A | 2 | 1h88C | 3 | 3r05A | 7 | 4il6B | 2dis |
| 3h2tA | 2 | 1m8pB | 3 | 3ubhA | 4 | 4jxkA | 2dis |
| 3hcsA | 2 | 1ni5A | 3 | 3w2wA | 4 | 4m8mB | 2dis |
| 3hyiA | 2 | 1q25A | 3 | 3zniA | 4 | 4mzyA | 2dis |
| 3i2dA | 2 | 1uzjA | 3 | 4aimA | 4 | 4onyA | 2dis |
| 3iam2 | 2 | 1zpuA | 3 | 4ak1A | 6 | 4pyhA | 2dis |
| 3ifrA | 2 | 1zy9A | 3 | 4aq1A | 6 | 4rg1A | 2dis |

**Table S4.** Detailed results of E2EDA, SADA, DEMO and AIDA using native structures for domain assembly on 356 benchmark proteins.

| PDB ID | TM-score | | | | PDB ID | TM-score | | | |
|---|---|---|---|---|---|---|---|---|---|
| | E2EDA | SADA | DEMO | AIDA | | E2EDA | SADA | DEMO | AIDA |
| 1cjyA | 0.82 | 0.81 | 0.81 | 0.81 | 2b5uA | 0.79 | 0.48 | 0.45 | 0.51 |
| 1efdN | 0.92 | 0.97 | 1 | 0.71 | 2ewfA | 0.5 | 0.49 | 0.55 | 0.47 |
| 1fjrA | 0.67 | 0.86 | 0.75 | 0.68 | 2piaA | 0.72 | 0.86 | 0.75 | 0.74 |
| 1g87B | 0.98 | 0.76 | 0.78 | 0.79 | 2r7dA | 0.98 | 0.98 | 0.92 | 0.75 |
| 1hx6B | 0.8 | 0.98 | 0.93 | 0.65 | 2uwnA | 0.8 | 0.77 | 0.48 | 0.37 |
| 1iwaA | 0.99 | 0.99 | 1 | 0.82 | 2v0nA | 0.41 | 0.45 | 0.41 | 0.41 |
| 1m5qH | 0.59 | 0.58 | 0.57 | 0.56 | 2vgmA | 0.67 | 0.76 | 0.66 | 0.4 |
| 1mkfA | 0.54 | 0.99 | 0.95 | 0.61 | 2wqrB | 0.73 | 0.49 | 0.67 | 0.37 |
| 1mkmB | 0.97 | 0.7 | 0.7 | 0.7 | 2y25B | 0.63 | 0.57 | 0.42 | 0.4 |
| 1nh2D | 0.92 | 0.58 | 0.54 | 0.53 | 2yk0A | 0.7 | 0.72 | 0.74 | 0.43 |
| 1pprM | 0.86 | 0.57 | 0.99 | 0.93 | 2zzqA | 0.97 | 0.46 | 0.72 | 0.72 |
| 1prrA | 0.53 | 0.6 | 0.53 | 0.55 | 3bt1U | 0.89 | 0.61 | 0.4 | 0.42 |
| 1q19A | 0.8 | 0.99 | 0.96 | 0.95 | 3c1yA | 0.99 | 0.63 | 0.49 | 0.49 |
| 1qwrA | 0.96 | 1 | 0.98 | 0.75 | 3cw2C | 0.58 | 0.7 | 0.65 | 0.66 |
| 1r71B | 0.84 | 0.88 | 0.73 | 0.55 | 3f83A | 0.97 | 0.46 | 0.39 | 0.41 |
| 1rh1A | 0.71 | 0.88 | 0.55 | 0.58 | 3fc3A | 0.44 | 0.54 | 0.47 | 0.45 |
| 1rktA | 0.9 | 0.97 | 0.87 | 0.79 | 3gbgA | 0.98 | 0.64 | 0.6 | 0.63 |
| 1s6lA | 0.8 | 0.76 | 0.73 | 0.8 | 3h5cB | 0.91 | 0.97 | 0.86 | 0.6 |
| 1sp3A | 0.7 | 1 | 0.59 | 0.59 | 3ibjA | 0.5 | 0.58 | 0.5 | 0.49 |
| 1vz6A | 0.98 | 0.95 | 0.82 | 0.75 | 3ippB | 0.94 | 0.74 | 0.44 | 0.55 |
| 1w3aA | 0.59 | 0.62 | 0.57 | 0.54 | 3jymB | 0.92 | 0.43 | 0.49 | 0.44 |
| 1wv3A | 0.83 | 0.93 | 0.84 | 0.85 | 3kbgA | 0.86 | 0.62 | 0.76 | 0.46 |
| 1x7pA | 0.94 | 0.99 | 0.78 | 0.63 | 3mc8A | 0.68 | 0.83 | 0.87 | 0.51 |
| 1x9yA | 0.84 | 0.63 | 0.98 | 0.57 | 3npfA | 0.84 | 0.93 | 0.96 | 0.54 |
| 1y11A | 0.61 | 0.6 | 0.55 | 0.57 | 3orjA | 0.76 | 0.66 | 0.62 | 0.57 |
| 1yiqA | 0.99 | 0.88 | 0.99 | 0.89 | 3plaA | 0.66 | 0.65 | 0.63 | 0.63 |
| 1zbuB | 0.96 | 0.79 | 0.79 | 0.76 | 3qe9Y | 0.97 | 0.94 | 0.93 | 0.67 |
| 1ze1A | 0.94 | 0.99 | 0.96 | 0.79 | 3qjoA | 0.79 | 0.8 | 0.99 | 0.76 |
| 2ablA | 0.63 | 0.7 | 0.63 | 0.64 | 3qphA | 0.71 | 0.48 | 0.57 | 0.58 |
| 2ahvA | 0.97 | 0.82 | 0.75 | 0.8 | 3qyeA | 0.97 | 0.99 | 0.95 | 0.53 |
| 2bkpA | 0.6 | 0.83 | 0.59 | 0.79 | 3rimA | 0.96 | 0.98 | 0.97 | 0.54 |
| 2c1yA | 0.7 | 0.56 | 0.7 | 0.58 | 3rrpA | 0.95 | 0.98 | 0.97 | 0.66 |
| 2cxcA | 0.89 | 0.96 | 0.9 | 0.51 | 3soaA | 0.67 | 0.73 | 0.73 | 0.53 |
| 2d1cA | 0.94 | 0.84 | 0.79 | 0.78 | 3tixD | 0.94 | 0.63 | 0.61 | 0.39 |
| 2d7iA | 0.77 | 0.74 | 0.77 | 0.84 | 3tp9A | 0.55 | 0.87 | 0.57 | 0.59 |
| 2e9hA | 0.92 | 0.97 | 0.96 | 0.81 | 3ua3A | 0.57 | 0.58 | 0.51 | 0.45 |
| 2e9xB | 0.9 | 0.99 | 0.86 | 0.88 | 3uj0A | 0.95 | 0.94 | 0.56 | 0.5 |
| 2evrA | 0.9 | 0.99 | 0.69 | 0.68 | 3vn4A | 0.97 | 0.47 | 0.44 | 0.46 |
| 2ew9A | 0.56 | 0.54 | 0.57 | 0.57 | 3vsmA | 0.98 | 0.8 | 0.98 | 0.56 |
| 2fd5A | 0.97 | 0.99 | 0.97 | 0.75 | 3w1bA | 0.77 | 0.72 | 0.72 | 0.59 |
| 2gh8A | 0.9 | 0.72 | 0.66 | 0.78 | 3zh9B | 0.72 | 0.62 | 0.58 | 0.58 |
| 2gt1A | 0.95 | 0.97 | 0.96 | 0.64 | 4alzA | 0.6 | 0.69 | 0.38 | 0.46 |
| 2gzaC | 0.66 | 0.98 | 0.92 | 0.67 | 4ax8A | 0.69 | 0.64 | 0.69 | 0.52 |
| 2hjqA | 0.82 | 0.55 | 0.58 | 0.53 | 4b3iA | 0.85 | 0.94 | 0.68 | 0.5 |
| 2hwjA | 0.68 | 0.83 | 0.68 | 0.7 | 4bd9B | 0.67 | 0.61 | 0.51 | 0.39 |
| 2ijd1 | 0.74 | 0.77 | 0.75 | 0.76 | 4c0aB | 0.92 | 0.57 | 0.52 | 0.49 |
| 2iu7A | 0.93 | 0.6 | 0.58 | 0.58 | 4c0sA | 0.61 | 0.63 | 0.64 | 0.62 |
| 2iw2A | 0.99 | 0.98 | 0.96 | 0.67 | 4dimA | 0.76 | 0.89 | 0.9 | 0.62 |
| 2jz4A | 0.55 | 0.59 | 0.51 | 0.51 | 4indA | 0.45 | 0.46 | 0.42 | 0.41 |

| PDB ID | TM-score | | | | PDB ID | TM-score | | | |
|---|---|---|---|---|---|---|---|---|---|
| | E2EDA | SADA | DEMO | AIDA | | E2EDA | SADA | DEMO | AIDA |
| 2kdyA | 0.96 | 0.59 | 0.84 | 0.62 | 4jdzB | 0.93 | 0.7 | 0.63 | 0.45 |
| 2kn4A | 0.61 | 0.65 | 0.61 | 0.6 | 4kc3B | 0.66 | 0.76 | 0.67 | 0.37 |
| 2mbgA | 0.78 | 0.78 | 0.82 | 0.76 | 4kikB | 0.82 | 0.73 | 0.85 | 0.67 |
| 2nsfA | 0.89 | 0.94 | 0.72 | 0.68 | 4lmfA | 0.75 | 0.49 | 0.47 | 0.44 |
| 2nykA | 0.95 | 0.95 | 0.81 | 0.74 | 4lziA | 0.47 | 0.67 | 0.46 | 0.42 |
| 2o6yA | 0.96 | 0.99 | 0.98 | 0.67 | 4m9pA | 0.95 | 0.51 | 0.36 | 0.38 |
| 2owbA | 0.98 | 0.98 | 1 | 0.7 | 4pt5A | 0.91 | 0.91 | 0.82 | 0.46 |
| 2qfiA | 0.99 | 0.87 | 0.72 | 0.72 | 4uwhA | 0.95 | 0.99 | 0.94 | 0.49 |
| 2qp2A | 0.97 | 0.74 | 0.73 | 0.75 | 1c1zA | 0.75 | 0.46 | 0.41 | 0.3 |
| 2qygA | 0.99 | 0.97 | 0.99 | 0.68 | 1d2pA | 0.3 | 0.36 | 0.29 | 0.3 |
| 2r5wB | 0.97 | 0.8 | 0.6 | 0.58 | 1k7tA | 0.87 | 0.67 | 0.38 | 0.39 |
| 2uu7A | 0.98 | 0.99 | 0.96 | 0.75 | 1kfqA | 0.92 | 0.97 | 0.97 | 0.68 |
| 2w4bA | 0.98 | 0.83 | 0.79 | 0.77 | 1ldjA | 0.76 | 0.62 | 0.55 | 0.28 |
| 2x7iA | 0.89 | 0.98 | 0.96 | 0.58 | 1nyqB | 0.91 | 0.87 | 0.76 | 0.58 |
| 2x8kC | 0.92 | 0.98 | 0.71 | 0.83 | 1ug9A | 0.64 | 0.65 | 0.65 | 0.58 |
| 2yilA | 0.62 | 0.63 | 0.63 | 0.64 | 1z1wA | 0.96 | 0.93 | 0.95 | 0.5 |
| 2yrqA | 0.52 | 0.52 | 0.51 | 0.53 | 2au3A | 0.71 | 0.81 | 0.66 | 0.41 |
| 2zxcA | 0.99 | 0.82 | 0.81 | 0.8 | 2ii2A | 0.75 | 0.87 | 0.78 | 0.66 |
| 3a1iA | 0.9 | 0.93 | 0.9 | 0.89 | 2olsA | 0.46 | 0.49 | 0.5 | 0.44 |
| 3a45A | 0.98 | 0.99 | 0.99 | 0.54 | 2ra1A | 0.6 | 0.49 | 0.33 | 0.44 |
| 3a56A | 0.61 | 0.66 | 0.76 | 0.66 | 2v5dA | 0.75 | 0.69 | 0.82 | 0.59 |
| 3ajvA | 0.85 | 0.98 | 0.76 | 0.55 | 2xt6A | 0.98 | 0.54 | 0.41 | 0.38 |
| 3aqkA | 0.95 | 0.96 | 0.96 | 0.73 | 2zpaB | 0.94 | 0.9 | 0.44 | 0.41 |
| 3arbA | 0.96 | 0.94 | 0.95 | 0.85 | 3apoA | 0.7 | 0.45 | 0.41 | 0.29 |
| 3aujG | 0.98 | 0.99 | 0.7 | 0.71 | 3b43A | 0.36 | 0.36 | 0.44 | 0.32 |
| 3b2zF | 0.95 | 0.76 | 0.76 | 0.81 | 3gf5B | 0.53 | 0.41 | 0.46 | 0.31 |
| 3b7wA | 0.96 | 0.83 | 0.83 | 0.83 | 3hjlA | 0.36 | 0.36 | 0.39 | 0.38 |
| 3bt3A | 0.85 | 0.95 | 0.92 | 0.62 | 3kq4B | 0.39 | 0.36 | 0.35 | 0.41 |
| 3c4tA | 0.83 | 0.74 | 0.74 | 0.74 | 3kwlA | 0.47 | 0.58 | 0.53 | 0.38 |
| 3craA | 0.93 | 0.61 | 0.58 | 0.54 | 3ob8A | 0.95 | 0.97 | 0.68 | 0.47 |
| 3d30A | 0.89 | 0.96 | 0.91 | 0.57 | 3opfB | 0.77 | 0.68 | 0.63 | 0.42 |
| 3eo5A | 0.97 | 0.64 | 0.6 | 0.53 | 3p53A | 0.46 | 0.37 | 0.35 | 0.37 |
| 3errA | 0.8 | 0.79 | 0.64 | 0.66 | 3pvlA | 0.9 | 0.68 | 0.75 | 0.51 |
| 3g79A | 0.95 | 0.9 | 0.84 | 0.6 | 3r05A | 0.35 | 0.32 | 0.31 | 0.3 |
| 3h2tA | 0.98 | 0.81 | 0.82 | 0.86 | 3ubhA | 0.65 | 0.68 | 0.51 | 0.33 |
| 3hcsA | 0.82 | 0.8 | 0.7 | 0.69 | 3w2wA | 0.82 | 0.85 | 0.51 | 0.53 |
| 3hyiA | 0.69 | 0.71 | 0.69 | 0.68 | 3zniA | 0.79 | 0.41 | 0.35 | 0.4 |
| 3i2dA | 0.87 | 0.86 | 0.78 | 0.56 | 4aimA | 0.8 | 0.91 | 0.51 | 0.49 |
| 3iam2 | 0.99 | 0.92 | 0.64 | 0.6 | 4ak1A | 0.28 | 0.29 | 0.28 | 0.29 |
| 3ifrA | 0.96 | 0.99 | 0.98 | 0.65 | 4aq1A | 0.81 | 0.45 | 0.46 | 0.35 |
| 3isqA | 0.95 | 1 | 0.99 | 0.62 | 4fe9A | 0.91 | 0.47 | 0.44 | 0.38 |
| 3j7aK | 0.94 | 0.96 | 0.99 | 0.71 | 4h2aA | 0.77 | 0.61 | 0.53 | 0.43 |
| 3k1rA | 0.62 | 0.66 | 0.61 | 0.58 | 4i5sB | 0.5 | 0.55 | 0.4 | 0.41 |
| 3k2iA | 0.93 | 0.78 | 0.73 | 0.7 | 4iggB | 0.44 | 0.31 | 0.33 | 0.31 |
| 3kh5A | 0.95 | 0.94 | 0.98 | 0.56 | 4j9vA | 0.96 | 0.87 | 0.61 | 0.46 |
| 3kjpA | 0.86 | 0.91 | 0.71 | 0.53 | 4k3bA | 0.6 | 0.56 | 0.54 | 0.49 |
| 3kt1A | 0.97 | 0.91 | 0.58 | 0.58 | 4kwuA | 0.96 | 0.78 | 0.82 | 0.46 |
| 3ktmE | 0.73 | 0.63 | 0.59 | 0.62 | 4m00A | 0.49 | 0.52 | 0.51 | 0.52 |
| 3kzwA | 1 | 1 | 0.99 | 0.68 | 1bp1A | 0.98 | 0.99 | 0.98 | 0.73 |
| 3l76A | 0.97 | 0.89 | 0.65 | 0.72 | 1cjsA | 0.94 | 0.84 | 0.97 | 0.64 |
| 3ld1A | 0.7 | 0.7 | 0.7 | 0.7 | 1ck1A | 0.99 | 0.99 | 0.95 | 0.64 |
| 3lsgA | 0.99 | 0.86 | 0.86 | 0.61 | 1ecrA | 1 | 0.88 | 0.87 | 0.85 |
| 3me4A | 0.96 | 0.9 | 0.63 | 0.6 | 1f5qD | 0.91 | 0.99 | 0.95 | 0.77 |

| PDB ID | TM-score | | | | PDB ID | TM-score | | | |
|---|---|---|---|---|---|---|---|---|---|
| | E2EDA | SADA | DEMO | AIDA | | E2EDA | SADA | DEMO | AIDA |
| 3ml4C | 0.58 | 0.65 | 0.93 | 0.55 | 1fa9A | 0.99 | 1 | 0.94 | 0.68 |
| 3mx2B | 0.97 | 0.92 | 0.89 | 0.73 | 1gu7A | 0.96 | 1 | 0.94 | 0.61 |
| 3mzfA | 0.91 | 0.93 | 0.86 | 0.75 | 1itwA | 0.97 | 0.99 | 0.61 | 0.62 |
| 3njaB | 0.59 | 0.61 | 0.61 | 0.58 | 1jkiA | 0.98 | 1 | 0.91 | 0.82 |
| 3nqiA | 0.93 | 0.94 | 0.93 | 0.74 | 1n80A | 0.96 | 1 | 0.84 | 0.78 |
| 3nt8A | 0.94 | 0.76 | 0.58 | 0.52 | 1nzjA | 0.97 | 0.99 | 0.98 | 0.82 |
| 3og5A | 0.9 | 0.97 | 0.92 | 0.78 | 1qhdA | 0.96 | 0.9 | 0.88 | 0.6 |
| 3oh0A | 0.94 | 0.97 | 0.78 | 0.83 | 1qz9A | 0.82 | 0.99 | 0.99 | 0.77 |
| 3pcsB | 0.99 | 0.86 | 0.8 | 0.76 | 1sb7B | 0.98 | 0.87 | 0.97 | 0.64 |
| 3po3S | 0.93 | 0.96 | 0.58 | 0.56 | 1vk1A | 0.85 | 0.85 | 0.62 | 0.62 |
| 3pxpA | 0.74 | 0.74 | 0.7 | 0.69 | 1vrmA | 0.94 | 0.99 | 0.9 | 0.71 |
| 3qavA | 0.94 | 0.98 | 0.91 | 0.93 | 1xvuA | 0.97 | 0.98 | 0.91 | 0.82 |
| 3qf4B | 0.9 | 0.93 | 0.74 | 0.62 | 1yy3A | 0.91 | 0.7 | 0.7 | 0.7 |
| 3qjjA | 0.93 | 0.97 | 0.88 | 0.57 | 1z87A | 0.54 | 0.67 | 0.54 | 0.52 |
| 3qtdA | 0.98 | 1 | 0.99 | 0.62 | 2a1sC | 0.92 | 0.86 | 0.82 | 0.81 |
| 3r6bA | 0.97 | 0.54 | 0.65 | 0.51 | 2a3lA | 0.97 | 0.97 | 0.99 | 0.86 |
| 3rh7A | 0.99 | 0.71 | 0.61 | 0.8 | 2bt1A | 0.97 | 0.99 | 0.93 | 0.66 |
| 3rwxA | 0.82 | 0.6 | 0.55 | 0.58 | 2bydA | 0.85 | 0.98 | 0.58 | 0.63 |
| 3sb4A | 0.99 | 0.61 | 0.57 | 0.56 | 2c43A | 0.96 | 0.93 | 0.57 | 0.63 |
| 3swjA | 0.92 | 0.71 | 0.71 | 0.73 | 2dfyC | 0.96 | 0.66 | 0.62 | 0.48 |
| 3t58B | 0.94 | 0.58 | 0.55 | 0.53 | 2dlaA | 0.97 | 1 | 0.97 | 0.93 |
| 3t7jA | 0.91 | 0.97 | 0.98 | 0.52 | 2g3pA | 0.97 | 0.63 | 0.87 | 0.57 |
| 3u07C | 0.92 | 0.88 | 0.8 | 0.72 | 2gg6A | 1 | 0.99 | 0.99 | 0.72 |
| 3u0oB | 0.94 | 0.99 | 0.99 | 0.57 | 2gsyE | 0.95 | 0.92 | 0.7 | 0.68 |
| 3u9gA | 0.75 | 0.77 | 0.73 | 0.71 | 2gzoA | 0.71 | 0.93 | 0.57 | 0.53 |
| 3ub1D | 0.99 | 0.62 | 0.58 | 0.56 | 2j2cA | 0.95 | 0.96 | 0.72 | 0.66 |
| 3uitD | 0.55 | 0.57 | 0.52 | 0.55 | 2kfwA | 0.89 | 0.78 | 0.77 | 0.73 |
| 3uo3A | 0.87 | 0.99 | 0.9 | 0.83 | 2l9yA | 0.74 | 0.69 | 0.74 | 0.72 |
| 3v7oB | 0.63 | 0.65 | 0.65 | 0.62 | 2ntyB | 0.98 | 0.82 | 0.71 | 0.72 |
| 3vr8B | 0.95 | 1 | 0.66 | 0.61 | 2r3vA | 0.97 | 1 | 0.96 | 0.71 |
| 3wkuA | 0.77 | 0.76 | 0.78 | 0.77 | 2r58A | 0.99 | 0.65 | 0.99 | 0.53 |
| 3zvmA | 0.52 | 0.59 | 0.55 | 0.52 | 2w4mA | 0.96 | 0.89 | 0.78 | 0.77 |
| 4acoA | 0.99 | 0.91 | 0.8 | 0.76 | 2x0cA | 0.67 | 0.66 | 0.63 | 0.58 |
| 4ap5A | 0.94 | 0.99 | 0.74 | 0.62 | 2y51A | 0.98 | 0.99 | 0.98 | 0.83 |
| 4axdA | 0.95 | 0.99 | 0.88 | 0.77 | 2yb0E | 0.86 | 0.97 | 0.89 | 0.83 |
| 4bfiB | 0.85 | 0.83 | 0.96 | 0.59 | 2z86C | 0.96 | 0.69 | 0.76 | 0.57 |
| 4bt9B | 0.73 | 0.61 | 0.63 | 0.61 | 3afoA | 0.91 | 0.67 | 0.78 | 0.63 |
| 4cczA | 0.87 | 0.96 | 0.61 | 0.93 | 3bu2A | 0.97 | 0.91 | 0.68 | 0.7 |
| 4d0nB | 0.89 | 0.91 | 0.93 | 0.69 | 3cvzA | 0.98 | 0.87 | 0.61 | 0.61 |
| 4d1iG | 0.98 | 0.96 | 0.79 | 0.72 | 3dupA | 0.95 | 0.98 | 0.83 | 0.68 |
| 4dj3A | 0.97 | 0.98 | 0.94 | 0.61 | 3eswA | 0.95 | 0.89 | 0.89 | 0.91 |
| 4dqaA | 0.84 | 0.64 | 0.63 | 0.62 | 3eukH | 0.7 | 0.92 | 0.91 | 0.88 |
| 4eo3A | 0.57 | 0.58 | 0.57 | 0.59 | 3fi7A | 0.95 | 0.99 | 0.99 | 0.75 |
| 4eogA | 0.99 | 0.98 | 0.68 | 0.66 | 3fvvA | 0.78 | 0.98 | 0.93 | 0.72 |
| 4etxA | 0.98 | 0.55 | 0.55 | 0.6 | 3gmsA | 0.96 | 1 | 0.99 | 0.93 |
| 4fguA | 0.97 | 0.92 | 0.69 | 0.69 | 3hzzB | 0.99 | 1 | 0.93 | 0.79 |
| 4fkcA | 0.95 | 0.94 | 0.94 | 0.73 | 3m1uA | 0.95 | 0.98 | 0.6 | 0.73 |
| 4fxkC | 0.65 | 0.97 | 0.96 | 0.54 | 3mw8A | 0.83 | 0.75 | 0.94 | 0.55 |
| 4gbyA | 0.99 | 1 | 0.99 | 1 | 3mwcA | 0.97 | 1 | 0.98 | 0.91 |
| 4ggmX | 0.98 | 0.61 | 0.57 | 0.74 | 3nsjA | 0.97 | 0.82 | 0.84 | 0.78 |
| 4gslA | 0.96 | 0.55 | 0.54 | 0.53 | 3ntkA | 0.95 | 0.98 | 0.97 | 0.61 |
| 4gyjA | 0.96 | 1 | 1 | 0.67 | 3oaaG | 0.95 | 0.98 | 0.98 | 0.56 |
| 4h3tA | 0.92 | 0.98 | 0.89 | 0.79 | 3ptyA | 0.93 | 0.75 | 0.73 | 0.74 |

| PDB ID | TM-score | | | | PDB ID | TM-score | | | |
|---|---|---|---|---|---|---|---|---|---|
| | E2EDA | SADA | DEMO | AIDA | | E2EDA | SADA | DEMO | AIDA |
| 4hmoA | 0.9 | 0.99 | 0.94 | 0.55 | 3rfyA | 0.81 | 0.95 | 0.73 | 0.76 |
| 4ie6A | 0.99 | 0.7 | 0.7 | 0.7 | 3seoB | 0.66 | 0.77 | 0.65 | 0.61 |
| 4l5gA | 0.84 | 0.97 | 0.67 | 0.64 | 3spgA | 0.94 | 0.98 | 0.86 | 0.66 |
| 4lpqA | 0.95 | 0.91 | 0.7 | 0.66 | 3u0kA | 0.85 | 0.61 | 0.61 | 0.6 |
| 4m8rA | 0.76 | 0.77 | 0.79 | 0.76 | 3vlaA | 0.89 | 1 | 0.98 | 0.86 |
| 4n06B | 0.95 | 0.99 | 0.9 | 0.81 | 3vstA | 0.96 | 1 | 0.98 | 0.83 |
| 4nj5A | 0.87 | 0.69 | 0.69 | 0.61 | 4aqfB | 0.83 | 0.88 | 0.92 | 0.84 |
| 4opaB | 0.64 | 0.71 | 0.68 | 0.67 | 4b21A | 0.95 | 0.99 | 0.93 | 0.8 |
| 4qkuB | 0.98 | 1 | 0.73 | 0.72 | 4dt4A | 0.84 | 0.98 | 0.96 | 0.69 |
| 4up9A | 0.95 | 0.96 | 0.98 | 0.97 | 4dtfA | 0.84 | 0.95 | 0.85 | 0.81 |
| 4w7sA | 0.71 | 0.64 | 0.63 | 0.64 | 4ewtA | 0.97 | 0.84 | 0.88 | 0.72 |
| 1bf2A | 0.98 | 0.99 | 0.99 | 0.71 | 4f23A | 0.92 | 1 | 0.67 | 0.73 |
| 1bhgA | 0.96 | 1 | 0.97 | 0.58 | 4fzbC | 0.86 | 0.99 | 0.92 | 0.74 |
| 1f7uA | 0.97 | 0.97 | 0.76 | 0.64 | 4g1pA | 0.96 | 0.93 | 0.96 | 0.77 |
| 1fx7A | 0.66 | 0.87 | 0.78 | 0.63 | 4gfqA | 0.8 | 0.76 | 0.66 | 0.62 |
| 1griA | 0.72 | 0.68 | 0.52 | 0.52 | 4hvzA | 0.98 | 0.75 | 0.55 | 0.59 |
| 1h88C | 0.72 | 0.71 | 0.62 | 0.46 | 4il6B | 0.77 | 0.73 | 0.71 | 0.7 |
| 1m8pB | 0.96 | 0.68 | 0.7 | 0.38 | 4jxkA | 0.97 | 0.98 | 0.97 | 0.76 |
| 1ni5A | 0.71 | 0.75 | 0.67 | 0.42 | 4m8mB | 0.85 | 0.87 | 0.8 | 0.8 |
| 1q25A | 0.7 | 0.49 | 0.44 | 0.41 | 4mzyA | 0.97 | 1 | 0.95 | 0.79 |
| 1uzjA | 0.77 | 0.78 | 0.52 | 0.49 | 4onyA | 0.95 | 0.81 | 0.82 | 0.59 |
| 1zpuA | 0.89 | 0.99 | 0.99 | 0.5 | 4pyhA | 0.96 | 0.77 | 0.93 | 0.78 |
| 1zy9A | 0.91 | 0.92 | 0.96 | 0.45 | 4rg1A | 0.85 | 0.77 | 0.85 | 0.65 |

**Table S5.** TM-score for each model generated by AlphaFold2 and E2EDA on 185 human proteins.

| PDB ID | TM-score | | PDB ID | TM-score | | PDB ID | TM-score | |
|---|---|---|---|---|---|---|---|---|
| | E2EDA | AlphaFold2 | | E2EDA | AlphaFold2 | | E2EDA | AlphaFold2 |
| 1b3uA | 0.64 | 0.64 | 3dw8 | 0.76 | 0.75 | 5u6g | 0.37 | 0.37 |
| 1cdlA | 0.58 | 0.34 | 3eiqA | 0.66 | 0.64 | 5vgrA | 0.72 | 0.7 |
| 1dt9A | 0.84 | 0.79 | 3eiqD | 0.64 | 0.64 | 5wp9 | 0.44 | 0.44 |
| 1griA | 0.82 | 0.47 | 3fhtA | 0.57 | 0.66 | 5wsv | 0.41 | 0.33 |
| 1j1dD | 0.55 | 0.54 | 3fvyA | 0.72 | 0.68 | 5ztfA | 0.73 | 0.65 |
| 1j72A | 0.33 | 0.33 | 3g0hA | 0.87 | 0.64 | 6b9eA | 0.79 | 0.72 |
| 1n7dA | 0.23 | 0.24 | 3ikm | 0.75 | 0.75 | 6c3oE | 0.74 | 0.75 |
| 1nmv | 0.66 | 0.67 | 3j0aA | 0.62 | 0.61 | 6cesC | 0.77 | 0.76 |
| 1oqyA | 0.23 | 0.23 | 3jd8A | 0.69 | 0.7 | 6crfA | 0.75 | 0.64 |
| 1q8kA | 0.56 | 0.52 | 3k6sA | 0.23 | 0.24 | 6cwy | 0.67 | 0.68 |
| 1s8oA | 0.99 | 0.69 | 3l2oB | 0.88 | 0.65 | 6d6q | 0.7 | 0.73 |
| 1st0A | 0.82 | 0.77 | 3l4gC | 0.55 | 0.62 | 6dafA | 0.41 | 0.35 |
| 1te6A | 0.79 | 0.8 | 3l8iA | 0.68 | 0.67 | 6e5qA | 0.37 | 0.37 |
| 1v4tA | 0.76 | 0.75 | 3o96A | 0.93 | 0.71 | 6e6lA | 0.37 | 0.37 |
| 1xa6A | 0.59 | 0.37 | 3s3jA | 0.66 | 0.66 | 6e7iA | 0.79 | 0.79 |
| 1xmm | 0.78 | 0.77 | 3vklA | 0.6 | 0.56 | 6erpC | 0.74 | 0.77 |
| 1xtiA | 0.62 | 0.62 | 3wk4 | 0.99 | 0.69 | 6fonA | 0.62 | 0.62 |
| 1ytqA | 0.33 | 0.33 | 4a0cA | 0.77 | 0.8 | 6h25J | 0.71 | 0.72 |
| 1zbhA | 0.79 | 0.79 | 4a5tS | 0.24 | 0.23 | 6hcsA | 0.49 | 0.33 |
| 1zd8A | 0.57 | 0.57 | 4acqA | 0.55 | 0.64 | 6hhg | 0.89 | 0.68 |
| 2ar7A | 0.75 | 0.75 | 4axxA | 0.8 | 0.76 | 6hhjA | 0.86 | 0.66 |
| 2b3oA | 0.8 | 0.62 | 4bitA | 0.15 | 0.15 | 6i7sA | 0.59 | 0.62 |
| 2c9yA | 0.74 | 0.74 | 4bsm | 0.78 | 0.79 | 6jjuA | 0.72 | 0.66 |
| 2dvjA | 0.46 | 0.48 | 4cooA | 0.76 | 0.75 | 6jt0A | 0.45 | 0.55 |
| 2dybA | 0.88 | 0.7 | 4ct4B | 0.76 | 0.75 | 6jt0B | 0.49 | 0.45 |
| 2eyzA | 0.5 | 0.33 | 4d1eA | 0.37 | 0.42 | 6kvg | 0.63 | 0.48 |
| 2ffuA | 0.79 | 0.79 | 4d5nA | 0.72 | 0.67 | 6lyyA | 0.91 | 0.8 |
| 2gf5A | 0.79 | 0.48 | 4djcA | 0.59 | 0.59 | 6mue | 0.63 | 0.61 |
| 2havA | 0.22 | 0.22 | 4e0sA | 0.53 | 0.56 | 6nr8A | 0.68 | 0.69 |
| 2hxyA | 0.54 | 0.54 | 4ejnA | 0.74 | 0.72 | 6nu2z | 0.65 | 0.72 |
| 2j0uA | 0.56 | 0.56 | 4gow | 0.53 | 0.32 | 6nuiA | 0.73 | 0.74 |
| 2jdfA | 0.62 | 0.62 | 4h1yP | 0.69 | 0.69 | 6o81 | 0.74 | 0.75 |
| 2k0jA | 0.41 | 0.34 | 4h2fA | 0.67 | 0.67 | 6r6hB | 0.61 | 0.72 |
| 2kdoA | 0.41 | 0.28 | 4hanA | 0.54 | 0.57 | 6r7iG | 0.8 | 0.79 |
| 2kn6A | 0.52 | 0.46 | 4iggA | 0.57 | 0.65 | 6s7pA | 0.69 | 0.69 |
| 2kr0A | 0.48 | 0.3 | 4iggB | 0.53 | 0.66 | 6sn1A | 0.68 | 0.69 |
| 2l4hA | 0.75 | 0.72 | 4kfzA | 0.49 | 0.48 | 6tgbB | 0.61 | 0.7 |
| 2l6lA | 0.39 | 0.38 | 4ktyA | 0.6 | 0.6 | 6tntN | 0.62 | 0.61 |
| 2l9nA | 0.63 | 0.48 | 4l0dA | 0.76 | 0.75 | 6u3aA | 0.54 | 0.34 |
| 2looA | 0.35 | 0.32 | 4l27B | 0.94 | 0.75 | 6ulgF | 0.8 | 0.77 |

| PDB ID | TM-score | | PDB ID | TM-score | | PDB ID | TM-score | |
|---|---|---|---|---|---|---|---|---|
| | E2EDA | AlphaFold2 | | E2EDA | AlphaFold2 | | E2EDA | AlphaFold2 |
| 2lqlA | 0.37 | 0.38 | 4ll4C | 0.54 | 0.5 | 6ulgL | 0.58 | 0.52 |
| 2lqnA | 0.49 | 0.36 | 4nqaB | 0.71 | 0.59 | 6umx | 0.69 | 0.72 |
| 2lv6A | 0.39 | 0.34 | 4qdeA | 0.96 | 0.79 | 6wm2 | 0.75 | 0.69 |
| 2mph | 0.7 | 0.5 | 4rr2D | 0.61 | 0.61 | 6xteA | 0.28 | 0.27 |
| 2nn6G | 0.62 | 0.6 | 4ybhA | 0.73 | 0.76 | 6z3w | 0.79 | 0.8 |
| 2nn6I | 0.8 | 0.75 | 5a3fC | 0.72 | 0.64 | 7a09J | 0.77 | 0.78 |
| 2octA | 0.47 | 0.47 | 5a8lQ | 0.69 | 0.73 | 7abio | 0.64 | 0.64 |
| 2ok5A | 0.19 | 0.21 | 5c19A | 0.93 | 0.77 | 7apkG | 0.45 | 0.45 |
| 2pkgA | 0.78 | 0.74 | 5dow | 0.69 | 0.52 | 7azpA | 0.2 | 0.19 |
| 2q3zA | 0.67 | 0.67 | 5edm | 0.19 | 0.18 | 7bp9 | 0.8 | 0.79 |
| 2qnaA | 0.75 | 0.72 | 5ftnA | 0.75 | 0.7 | 7c3m | 0.86 | 0.7 |
| 2rsvA | 0.73 | 0.74 | 5gjqJ | 0.66 | 0.64 | 7cccB | 0.64 | 0.61 |
| 2wzb | 0.7 | 0.66 | 5gjqK | 0.84 | 0.78 | 7cunI | 0.76 | 0.75 |
| 2xw9 | 0.18 | 0.17 | 5ivw | 0.74 | 0.74 | 7dme | 0.44 | 0.45 |
| 2y3iA | 0.75 | 0.65 | 5js7A | 0.67 | 0.67 | 7egb4 | 0.76 | 0.78 |
| 2y4tA | 0.79 | 0.79 | 5jthA | 0.32 | 0.32 | 7jtnA | 0.22 | 0.21 |
| 2ymb | 0.95 | 0.68 | 5kcvA | 0.94 | 0.74 | 7kzpA | 0.58 | 0.6 |
| 2yrqA | 0.24 | 0.25 | 5nx2A | 0.74 | 0.76 | 7kzvV | 0.7 | 0.73 |
| 2zu6A | 0.68 | 0.58 | 5oeoA | 0.27 | 0.27 | 7lbmr | 0.58 | 0.58 |
| 3ajmA | 0.67 | 0.67 | 5t0gC | 0.6 | 0.6 | 7nfyA | 0.63 | 0.56 |
| 3ba0A | 0.52 | 0.53 | 5t7cA | 0.37 | 0.37 | 7nvr2 | 0.8 | 0.79 |
| 3bl7A | 0.85 | 0.8 | 5tp6A | 0.5 | 0.3 | | | |

**Table S6.** Detailed results for each target protein of E2EDA and AlphaFold2 on the multi-domain protein of CASP14.

| PBD ID | TM-score | | | PBD ID | TM-score | | |
|---|---|---|---|---|---|---|---|
| | E2EDA | E2EDA-native | AlphaFold2 | | E2EDA | E2EDA-native | AlphaFold2 |
| T1024 | 0.95 | 1 | 0.71 | T1070 | 0.62 | 0.75 | 0.47 |
| T1030 | 0.79 | 0.87 | 0.73 | T1091 | 0.7 | 0.78 | 0.83 |
| T1038 | 0.92 | 0.99 | 0.91 | T1092 | 0.92 | 0.94 | 0.92 |
| T1047s2 | 0.79 | 0.83 | 0.79 | T1093 | 0.86 | 0.89 | 0.94 |
| T1050 | 0.93 | 0.95 | 0.98 | T1094 | 0.91 | 0.99 | 0.9 |
| T1052 | 0.7 | 0.71 | 0.7 | T1096 | 0.55 | 0.58 | 0.55 |
| T1053 | 0.93 | 0.97 | 0.98 | T1100 | 0.94 | 0.88 | 0.93 |
| T1058 | 0.95 | 0.95 | 0.96 | T1101 | 0.89 | 0.94 | 0.93 |
| T1061 | 0.74 | 0.83 | 0.75 | | | | |

**Table S7.** The average running time of E2EDA and SADA for different types of proteins in the benchmark protein.

| Method | Average runtime (h) | | | | | *P*-value |
|---|---|---|---|---|---|---|
| | 2dis | 2dom | 3dom | m4dom | Total | |
| **E2EDA** | **0.50** | **0.57** | **0.73** | **0.91** | **0.62** | NA |
| SADA | 2.33 | 2.15 | 2.53 | 3.75 | 2.44 | 9.20E-67 |

**Table S8.** Detailed run times for each protein of E2EDA and SADA on 356 benchmark proteins.

| PDB ID | Length | TM-score | | PDB ID | Length | TM-score | |
|---|---|---|---|---|---|---|---|
| | | E2EDA | SADA | | | E2EDA | SADA |
| 1bp1A | 456 | 0.46 | 3.02 | 3kzwA | 494 | 0.5 | 3.97 |
| 1cjsA | 212 | 0.35 | 0.90 | 3l76A | 585 | 0.6 | 4.94 |
| 1ck1A | 239 | 0.3 | 1.27 | 3ld1A | 359 | 0.35 | 2.50 |
| 1ecrA | 305 | 0.31 | 1.84 | 3lsgA | 101 | 0.6 | 0.14 |
| 1f5qD | 248 | 0.35 | 1.34 | 3me4A | 169 | 0.3 | 0.53 |
| 1fa9A | 833 | 0.81 | 8.50 | 3ml4C | 203 | 0.27 | 0.76 |
| 1gu7A | 364 | 1.39 | 2.23 | 3mx2B | 517 | 0.39 | 4.34 |
| 1itwA | 740 | 0.6 | 6.64 | 3mzfA | 362 | 0.57 | 2.23 |
| 1jkiA | 525 | 0.43 | 4.19 | 3njaB | 107 | 0.94 | 0.14 |
| 1n80A | 328 | 0.29 | 1.93 | 3nqiA | 240 | 0.3 | 1.05 |
| 1nzjA | 273 | 0.34 | 1.54 | 3nt8A | 401 | 0.79 | 2.87 |
| 1qhdA | 397 | 0.32 | 2.66 | 3og5A | 155 | 0.46 | 0.36 |
| 1qz9A | 404 | 1.17 | 2.76 | 3oh0A | 593 | 0.61 | 5.63 |
| 1sb7B | 340 | 0.33 | 2.18 | 3pcsB | 355 | 0.41 | 2.36 |
| 1vk1A | 223 | 0.36 | 0.97 | 3po3S | 164 | 0.33 | 0.49 |
| 1vrmA | 304 | 0.34 | 1.83 | 3pxpA | 288 | 0.62 | 1.47 |
| 1xvuA | 402 | 0.58 | 2.80 | 3qavA | 219 | 0.55 | 1.07 |
| 1yy3A | 315 | 0.33 | 1.87 | 3qf4B | 588 | 3.33 | 4.70 |
| 1z87A | 263 | 0.53 | 1.39 | 3qjjA | 243 | 0.36 | 1.27 |
| 2a1sC | 393 | 0.35 | 2.81 | 3qtdA | 435 | 0.55 | 2.86 |
| 2a3lA | 616 | 0.53 | 5.98 | 3r6bA | 115 | 0.43 | 0.21 |
| 2bt1A | 254 | 0.33 | 1.40 | 3rh7A | 289 | 0.73 | 1.61 |
| 2bydA | 283 | 0.38 | 1.52 | 3rwxA | 239 | 0.33 | 1.12 |
| 2c43A | 280 | 0.36 | 1.47 | 3sb4A | 323 | 0.69 | 2.01 |
| 2dfyC | 158 | 0.37 | 0.45 | 3swjA | 246 | 0.45 | 1.21 |
| 2dlaA | 222 | 0.35 | 0.99 | 3t58B | 509 | 0.8 | 4.16 |
| 2g3pA | 201 | 0.38 | 0.82 | 3t7jA | 237 | 0.34 | 1.27 |
| 2gg6A | 445 | 0.52 | 3.30 | 3u07C | 382 | 0.42 | 2.47 |
| 2gsyE | 433 | 0.4 | 3.10 | 3u0oB | 335 | 0.47 | 2.02 |
| 2gzoA | 187 | 0.34 | 0.70 | 3u9gA | 217 | 0.3 | 1.10 |
| 2j2cA | 470 | 0.45 | 3.65 | 3ub1D | 251 | 0.33 | 1.30 |
| 2kfwA | 196 | 0.44 | 0.85 | 3uitD | 257 | 0.33 | 1.36 |
| 2l9yA | 167 | 0.54 | 0.52 | 3uo3A | 184 | 0.46 | 0.59 |
| 2ntyB | 337 | 0.35 | 2.15 | 3v7oB | 203 | 0.36 | 0.86 |
| 2r3vA | 390 | 0.48 | 2.95 | 3vr8B | 249 | 0.5 | 1.42 |
| 2r58A | 212 | 0.34 | 0.96 | 3wkuA | 405 | 0.47 | 2.97 |
| 2w4mA | 250 | 0.61 | 1.26 | 3zvmA | 385 | 1.42 | 2.65 |
| 2x0cA | 308 | 0.37 | 1.79 | 4acoA | 452 | 0.49 | 3.64 |
| 2y51A | 529 | 1.03 | 4.12 | 4ap5A | 373 | 0.39 | 2.56 |
| 2yb0E | 264 | 0.34 | 1.49 | 4axdA | 437 | 0.41 | 2.91 |

| PDB ID | Length | TM-score | | PDB ID | Length | TM-score | |
|---|---|---|---|---|---|---|---|
| | | E2EDA | SADA | | | E2EDA | SADA |
| 2z86C | 603 | 0.94 | 5.18 | 4bfiB | 202 | 0.53 | 0.85 |
| 3afoA | 360 | 0.42 | 2.23 | 4bt9B | 228 | 1.25 | 1.05 |
| 3bu2A | 189 | 0.31 | 0.66 | 4cczA | 611 | 0.62 | 5.41 |
| 3cvzA | 248 | 0.28 | 1.28 | 4d0nB | 371 | 0.43 | 2.56 |
| 3dupA | 279 | 0.44 | 1.51 | 4d1iG | 544 | 0.63 | 4.49 |
| 3eswA | 332 | 0.34 | 2.03 | 4dj3A | 298 | 2.02 | 1.73 |
| 3eukH | 459 | 1.2 | 3.40 | 4dqaA | 358 | 1.21 | 2.07 |
| 3fi7A | 177 | 0.28 | 0.63 | 4eo3A | 328 | 0.95 | 2.00 |
| 3fvvA | 223 | 0.52 | 0.99 | 4eogA | 460 | 0.46 | 3.33 |
| 3gmsA | 331 | 0.84 | 1.98 | 4etxA | 300 | 1.87 | 1.63 |
| 3hzzB | 447 | 0.89 | 3.39 | 4fguA | 407 | 0.38 | 2.90 |
| 3m1uA | 410 | 0.41 | 2.79 | 4fkcA | 370 | 0.51 | 2.53 |
| 3mw8A | 233 | 0.32 | 0.99 | 4fxkC | 290 | 0.33 | 1.61 |
| 3mwcA | 388 | 0.48 | 2.63 | 4gbyA | 475 | 2.2 | 3.40 |
| 3nsjA | 530 | 0.5 | 4.51 | 4ggmX | 270 | 0.31 | 1.40 |
| 3ntkA | 169 | 0.29 | 0.48 | 4gslA | 598 | 0.75 | 4.89 |
| 3oaaG | 284 | 0.32 | 1.53 | 4gyjA | 618 | 0.72 | 5.38 |
| 3ptyA | 284 | 0.29 | 1.47 | 4h3tA | 518 | 0.4 | 4.24 |
| 3rfyA | 356 | 0.45 | 2.05 | 4hmoA | 310 | 0.45 | 1.72 |
| 3seoB | 226 | 0.28 | 0.99 | 4ie6A | 404 | 0.41 | 2.78 |
| 3spgA | 332 | 0.48 | 2.04 | 4l5gA | 162 | 0.31 | 0.40 |
| 3u0kA | 396 | 1.54 | 2.58 | 4lpqA | 206 | 0.55 | 0.73 |
| 3vlaA | 428 | 0.45 | 2.95 | 4m8rA | 398 | 0.34 | 2.91 |
| 3vstA | 638 | 1.25 | 6.65 | 4n06B | 340 | 0.34 | 2.07 |
| 4aqfB | 474 | 0.39 | 3.67 | 4nj5A | 482 | 0.52 | 3.46 |
| 4b21A | 207 | 0.35 | 0.88 | 4opaB | 197 | 0.32 | 0.77 |
| 4dt4A | 160 | 0.33 | 0.41 | 4qkuB | 427 | 0.47 | 3.15 |
| 4dtfA | 359 | 0.32 | 2.18 | 4up9A | 514 | 0.63 | 4.22 |
| 4ewtA | 389 | 0.7 | 2.58 | 4w7sA | 444 | 1.17 | 3.15 |
| 4f23A | 509 | 0.44 | 3.58 | 1bf2A | 750 | 0.82 | 6.87 |
| 4fzbC | 207 | 0.31 | 0.95 | 1bhgA | 611 | 0.93 | 4.25 |
| 4g1pA | 479 | 0.62 | 3.57 | 1f7uA | 606 | 0.64 | 4.79 |
| 4gfqA | 186 | 0.32 | 0.57 | 1fx7A | 230 | 0.79 | 0.77 |
| 4hvzA | 214 | 0.3 | 0.83 | 1griA | 211 | 0.5 | 0.72 |
| 4il6B | 505 | 0.45 | 3.75 | 1h88C | 152 | 0.43 | 0.31 |
| 4jxkA | 327 | 0.96 | 1.98 | 1m8pB | 571 | 0.94 | 3.87 |
| 4m8mB | 574 | 0.45 | 4.87 | 1ni5A | 428 | 0.65 | 2.68 |
| 4mzyA | 495 | 0.43 | 3.59 | 1q25A | 428 | 0.4 | 2.83 |
| 4onyA | 583 | 1.05 | 4.74 | 1uzjA | 162 | 0.43 | 0.37 |
| 4pyhA | 293 | 0.4 | 1.60 | 1zpuA | 504 | 0.62 | 3.52 |
| 4rg1A | 285 | 0.28 | 1.53 | 1zy9A | 522 | 0.59 | 3.77 |
| 1cjyA | 630 | 0.71 | 6.20 | 2b5uA | 465 | 0.48 | 2.65 |
| 1efdN | 262 | 0.41 | 1.18 | 2ewfA | 574 | 0.51 | 3.88 |

| PDB ID | Length | TM-score | | PDB ID | Length | TM-score | |
|---|---|---|---|---|---|---|---|
| | | E2EDA | SADA | | | E2EDA | SADA |
| 1fjrA | 188 | 0.35 | 0.67 | 2piaA | 321 | 0.59 | 1.70 |
| 1g87B | 613 | 0.6 | 5.58 | 2r7dA | 452 | 0.49 | 3.09 |
| 1hx6B | 396 | 0.42 | 2.54 | 2uwnA | 187 | 0.34 | 0.56 |
| 1iwaA | 441 | 0.42 | 3.46 | 2v0nA | 459 | 2.61 | 3.04 |
| 1m5qH | 123 | 0.36 | 0.21 | 2vgmA | 354 | 0.44 | 2.07 |
| 1mkfA | 371 | 0.32 | 2.47 | 2wqrB | 322 | 0.87 | 1.63 |
| 1mkmB | 247 | 0.74 | 1.30 | 2y25B | 314 | 0.9 | 1.57 |
| 1nh2D | 102 | 0.29 | 0.14 | 2yk0A | 698 | 0.65 | 5.06 |
| 1pprM | 312 | 0.33 | 1.74 | 2zzqA | 469 | 0.47 | 3.18 |
| 1prrA | 173 | 0.33 | 0.51 | 3bt1U | 273 | 0.29 | 1.23 |
| 1q19A | 500 | 0.77 | 3.87 | 3c1yA | 349 | 0.41 | 2.05 |
| 1qwrA | 321 | 0.37 | 1.88 | 3cw2C | 251 | 0.44 | 0.98 |
| 1r71B | 116 | 0.36 | 0.16 | 3f83A | 508 | 0.46 | 3.25 |
| 1rh1A | 490 | 0.45 | 3.47 | 3fc3A | 190 | 0.36 | 0.51 |
| 1rktA | 207 | 0.92 | 0.90 | 3gbgA | 260 | 2.01 | 1.16 |
| 1s6lA | 181 | 0.33 | 0.68 | 3h5cB | 263 | 0.74 | 1.22 |
| 1sp3A | 436 | 0.51 | 3.11 | 3ibjA | 661 | 2.29 | 4.97 |
| 1vz6A | 365 | 0.37 | 2.47 | 3ippB | 420 | 0.8 | 2.60 |
| 1w3aA | 312 | 0.35 | 1.92 | 3jymB | 307 | 0.45 | 1.53 |
| 1wv3A | 181 | 0.44 | 0.66 | 3kbgA | 187 | 0.39 | 0.47 |
| 1x7pA | 265 | 0.34 | 1.33 | 3mc8A | 257 | 0.57 | 1.05 |
| 1x9yA | 346 | 0.34 | 2.28 | 3npfA | 306 | 0.61 | 1.49 |
| 1y11A | 356 | 1.04 | 2.25 | 3orjA | 415 | 0.66 | 2.52 |
| 1yiqA | 684 | 2.2 | 7.91 | 3plaA | 375 | 0.41 | 2.20 |
| 1zbuB | 289 | 0.37 | 1.68 | 3qe9Y | 345 | 0.49 | 1.82 |
| 1ze1A | 308 | 0.32 | 1.86 | 3qjoA | 491 | 0.52 | 3.47 |
| 2ablA | 163 | 0.29 | 0.49 | 3qphA | 335 | 0.69 | 1.84 |
| 2ahvA | 512 | 0.45 | 3.95 | 3qyeA | 317 | 0.44 | 1.68 |
| 2bkpA | 202 | 0.3 | 0.70 | 3rimA | 697 | 1.02 | 5.67 |
| 2c1yA | 231 | 0.39 | 1.06 | 3rrpA | 455 | 0.55 | 2.91 |
| 2cxcA | 138 | 0.29 | 0.31 | 3soaA | 436 | 1.93 | 2.78 |
| 2d1cA | 495 | 0.49 | 4.36 | 3tixD | 424 | 0.38 | 2.78 |
| 2d7iA | 536 | 1.54 | 5.04 | 3tp9A | 467 | 0.84 | 3.07 |
| 2e9hA | 157 | 0.26 | 0.54 | 3ua3A | 648 | 1.15 | 4.73 |
| 2e9xB | 175 | 0.24 | 0.57 | 3uj0A | 502 | 0.63 | 3.22 |
| 2evrA | 240 | 0.37 | 0.98 | 3vn4A | 375 | 0.72 | 2.21 |
| 2ew9A | 149 | 0.34 | 0.35 | 3vsmA | 638 | 0.75 | 5.18 |
| 2fd5A | 180 | 0.56 | 0.52 | 3w1bA | 589 | 0.98 | 4.00 |
| 2gh8A | 544 | 0.41 | 4.19 | 3zh9B | 339 | 0.45 | 1.94 |
| 2gt1A | 323 | 0.39 | 1.79 | 4alzA | 212 | 0.44 | 0.50 |
| 2gzaC | 336 | 0.97 | 1.96 | 4ax8A | 456 | 1.7 | 2.88 |
| 2hjqA | 99 | 0.22 | 0.12 | 4b3iA | 736 | 1.51 | 5.81 |
| 2hwjA | 184 | 0.29 | 0.63 | 4bd9B | 165 | 0.37 | 0.40 |

| PDB ID | Length | TM-score | | PDB ID | Length | TM-score | |
|---|---|---|---|---|---|---|---|
| | | E2EDA | SADA | | | E2EDA | SADA |
| 2ijd1 | 644 | 0.6 | 6.43 | 4c0aB | 355 | 0.43 | 1.87 |
| 2iu7A | 159 | 0.35 | 0.37 | 4c0sA | 449 | 0.92 | 2.97 |
| 2iw2A | 479 | 0.68 | 3.79 | 4dimA | 382 | 1.05 | 2.23 |
| 2jz4A | 299 | 0.4 | 1.71 | 4indA | 383 | 0.39 | 2.32 |
| 2kdyA | 261 | 0.33 | 1.47 | 4jdzB | 447 | 0.81 | 2.65 |
| 2kn4A | 158 | 0.67 | 0.47 | 4kc3B | 278 | 1.01 | 1.25 |
| 2mbgA | 265 | 0.51 | 1.63 | 4kikB | 648 | 0.42 | 4.41 |
| 2nsfA | 240 | 0.42 | 1.02 | 4lmfA | 276 | 0.73 | 1.35 |
| 2nykA | 235 | 0.35 | 1.13 | 4lziA | 278 | 0.38 | 1.22 |
| 2o6yA | 514 | 0.53 | 4.03 | 4m9pA | 292 | 0.48 | 1.31 |
| 2owbA | 262 | 0.69 | 1.37 | 4pt5A | 319 | 0.39 | 1.76 |
| 2qfiA | 286 | 0.35 | 1.50 | 4uwhA | 553 | 0.92 | 3.68 |
| 2qp2A | 513 | 0.42 | 4.39 | 1c1zA | 325 | 0.44 | 1.25 |
| 2qygA | 429 | 0.47 | 3.30 | 1d2pA | 373 | 0.54 | 1.82 |
| 2r5wB | 345 | 0.97 | 2.42 | 1k7tA | 170 | 0.45 | 0.40 |
| 2uu7A | 370 | 0.44 | 2.41 | 1kfqA | 567 | 0.76 | 3.79 |
| 2w4bA | 455 | 0.39 | 3.75 | 1ldjA | 725 | 0.9 | 4.01 |
| 2x7iA | 296 | 0.39 | 1.72 | 1nyqB | 645 | 0.88 | 4.44 |
| 2x8kC | 243 | 0.38 | 1.21 | 1ug9A | 1019 | 1.41 | 8.63 |
| 2yilA | 131 | 0.41 | 0.31 | 1z1wA | 780 | 1.12 | 5.56 |
| 2yrqA | 173 | 0.44 | 0.54 | 2au3A | 403 | 0.97 | 2.04 |
| 2zxcA | 643 | 0.61 | 7.07 | 2ii2A | 304 | 0.43 | 1.21 |
| 3a1iA | 508 | 0.81 | 4.87 | 2olsA | 725 | 1.08 | 5.36 |
| 3a45A | 288 | 0.41 | 1.56 | 2ra1A | 413 | 0.56 | 1.84 |
| 3a56A | 291 | 0.36 | 1.46 | 2v5dA | 722 | 1.15 | 5.29 |
| 3ajvA | 174 | 0.37 | 0.44 | 2xt6A | 1055 | 1.53 | 8.69 |
| 3aqkA | 390 | 0.56 | 2.85 | 2zpaB | 651 | 1.55 | 4.41 |
| 3arbA | 274 | 0.45 | 1.46 | 3apoA | 688 | 1.64 | 3.52 |
| 3aujG | 139 | 0.34 | 0.32 | 3b43A | 569 | 0.49 | 2.79 |
| 3b2zF | 293 | 0.49 | 1.74 | 3gf5B | 380 | 0.48 | 1.29 |
| 3b7wA | 543 | 0.39 | 5.37 | 3hjlA | 321 | 0.47 | 1.37 |
| 3bt3A | 129 | 1.03 | 0.24 | 3kq4B | 457 | 0.67 | 2.71 |
| 3c4tA | 241 | 0.42 | 1.18 | 3kwlA | 493 | 0.5 | 2.94 |
| 3craA | 239 | 0.38 | 1.17 | 3ob8A | 1024 | 1.05 | 7.76 |
| 3d30A | 212 | 0.36 | 0.81 | 3opfB | 493 | 0.42 | 2.89 |
| 3eo5A | 169 | 0.4 | 0.44 | 3p53A | 494 | 0.44 | 2.91 |
| 3errA | 527 | 0.76 | 4.04 | 3pvlA | 598 | 0.61 | 3.32 |
| 3g79A | 475 | 0.84 | 3.54 | 3r05A | 1008 | 1.61 | 6.88 |
| 3h2tA | 326 | 0.37 | 2.07 | 3ubhA | 409 | 0.96 | 2.18 |
| 3hcsA | 157 | 0.59 | 0.52 | 3w2wA | 618 | 1.51 | 3.99 |
| 3hyiA | 286 | 0.06 | 1.65 | 3zniA | 391 | 0.59 | 1.85 |
| 3i2dA | 279 | 0.3 | 1.59 | 4aimA | 698 | 0.79 | 5.06 |
| 3iam2 | 179 | 0.28 | 0.60 | 4ak1A | 612 | 0.68 | 3.27 |

| PDB ID | Length | TM-score | | PDB ID | Length | TM-score | |
|---|---|---|---|---|---|---|---|
| | | E2EDA | SADA | | | E2EDA | SADA |
| 3ifrA | 483 | 0.54 | 3.50 | 4aq1A | 721 | 0.61 | 4.31 |
| 3isqA | 384 | 0.78 | 2.56 | 4fe9A | 450 | 0.36 | 2.38 |
| 3j7aK | 129 | 0.3 | 0.28 | 4h2aA | 708 | 0.77 | 5.08 |
| 3k1rA | 193 | 0.65 | 0.72 | 4i5sB | 395 | 2.84 | 2.02 |
| 3k2iA | 411 | 2.25 | 2.94 | 4iggB | 771 | 0.64 | 4.57 |
| 3kh5A | 279 | 1.06 | 1.74 | 4j9vA | 456 | 1.04 | 2.55 |
| 3kjpA | 293 | 0.32 | 1.97 | 4k3bA | 772 | 0.95 | 5.32 |
| 3kt1A | 557 | 0.5 | 4.60 | 4kwuA | 1030 | 1.47 | 7.59 |
| 3ktmE | 171 | 0.27 | 0.57 | 4m00A | 495 | 1.08 | 2.85 |

**Table S9.** Domain assembly results of E2EDA, E2EDA-w/o-S, and E2EDA-w/o-S&T on 356 benchmark proteins.

| PDB ID | TM-score | | | PDB ID | TM-score | | |
|---|---|---|---|---|---|---|---|
| | E2EDA | E2EDA -w/o-S | E2EDA -w/o-S&T | | E2EDA | E2EDA -w/o-S | E2EDA -w/o-S&T |
| 1cjyA | 0.82 | 0.84 | 0.86 | 3rwxA | 0.82 | 0.82 | 0.88 |
| 1efdN | 0.92 | 0.95 | 0.91 | 3sb4A | 0.99 | 0.98 | 0.92 |
| 1fjrA | 0.67 | 0.67 | 0.68 | 3swjA | 0.92 | 0.88 | 0.74 |
| 1g87B | 0.98 | 0.96 | 0.85 | 3t58B | 0.94 | 0.93 | 0.63 |
| 1hx6B | 0.8 | 0.8 | 0.71 | 3t7jA | 0.91 | 0.94 | 0.89 |
| 1iwaA | 0.99 | 0.99 | 0.8 | 3u07C | 0.92 | 0.92 | 0.8 |
| 1m5qH | 0.59 | 0.56 | 0.57 | 3u0oB | 0.94 | 0.96 | 0.97 |
| 1mkfA | 0.54 | 0.57 | 0.64 | 3u9gA | 0.75 | 0.74 | 0.85 |
| 1mkmB | 0.97 | 0.92 | 0.87 | 3ub1D | 0.99 | 0.95 | 0.58 |
| 1nh2D | 0.92 | 0.95 | 0.8 | 3uitD | 0.55 | 0.54 | 0.54 |
| 1pprM | 0.86 | 0.94 | 0.65 | 3uo3A | 0.87 | 0.87 | 0.88 |
| 1prrA | 0.53 | 0.53 | 0.53 | 3v7oB | 0.63 | 0.65 | 0.67 |
| 1q19A | 0.8 | 0.78 | 0.75 | 3vr8B | 0.95 | 0.97 | 0.79 |
| 1qwrA | 0.96 | 0.93 | 0.95 | 3wkuA | 0.77 | 0.78 | 0.86 |
| 1r71B | 0.84 | 0.75 | 0.75 | 3zvmA | 0.52 | 0.53 | 0.52 |
| 1rh1A | 0.71 | 0.71 | 0.84 | 4acoA | 0.99 | 0.94 | 0.77 |
| 1rktA | 0.9 | 0.93 | 0.89 | 4ap5A | 0.94 | 0.96 | 0.89 |
| 1s6lA | 0.8 | 0.79 | 0.73 | 4axdA | 0.95 | 0.92 | 0.79 |
| 1sp3A | 0.7 | 0.61 | 0.59 | 4bfiB | 0.85 | 0.92 | 0.92 |
| 1vz6A | 0.98 | 0.95 | 0.81 | 4bt9B | 0.73 | 0.58 | 0.66 |
| 1w3aA | 0.59 | 0.57 | 0.64 | 4cczA | 0.87 | 0.85 | 0.82 |
| 1wv3A | 0.83 | 0.73 | 0.69 | 4d0nB | 0.89 | 0.83 | 0.83 |
| 1x7pA | 0.94 | 0.95 | 0.8 | 4d1iG | 0.98 | 0.97 | 0.88 |
| 1x9yA | 0.84 | 0.61 | 0.65 | 4dj3A | 0.97 | 0.96 | 0.92 |
| 1y11A | 0.61 | 0.55 | 0.55 | 4dqaA | 0.84 | 0.6 | 0.71 |
| 1yiqA | 0.99 | 0.98 | 0.96 | 4eo3A | 0.57 | 0.57 | 0.57 |
| 1zbuB | 0.96 | 0.77 | 0.79 | 4eogA | 0.99 | 0.98 | 0.95 |
| 1ze1A | 0.94 | 0.99 | 0.98 | 4etxA | 0.98 | 0.55 | 0.55 |
| 2ablA | 0.63 | 0.63 | 0.63 | 4fguA | 0.97 | 0.96 | 0.73 |
| 2ahvA | 0.97 | 0.96 | 0.55 | 4fkcA | 0.95 | 0.99 | 0.95 |
| 2bkpA | 0.6 | 0.58 | 0.62 | 4fxkC | 0.65 | 0.6 | 0.61 |
| 2c1yA | 0.7 | 0.62 | 0.61 | 4gbyA | 0.99 | 0.98 | 0.98 |
| 2cxcA | 0.89 | 0.79 | 0.79 | 4ggmX | 0.98 | 0.91 | 0.94 |
| 2d1cA | 0.94 | 0.9 | 1 | 4gslA | 0.96 | 1 | 0.98 |
| 2d7iA | 0.77 | 0.77 | 0.87 | 4gyjA | 0.96 | 0.99 | 0.94 |
| 2e9hA | 0.92 | 0.88 | 0.94 | 4h3tA | 0.92 | 0.95 | 0.96 |
| 2e9xB | 0.9 | 0.86 | 0.79 | 4hmoA | 0.9 | 0.91 | 0.84 |
| 2evrA | 0.9 | 0.85 | 0.73 | 4ie6A | 0.99 | 0.98 | 0.69 |
| 2ew9A | 0.56 | 0.56 | 0.54 | 4l5gA | 0.84 | 0.64 | 0.65 |
| 2fd5A | 0.97 | 0.91 | 0.9 | 4lpqA | 0.95 | 0.94 | 0.85 |
| 2gh8A | 0.9 | 0.94 | 0.85 | 4m8rA | 0.76 | 0.78 | 0.86 |
| 2gt1A | 0.95 | 0.91 | 0.54 | 4n06B | 0.95 | 0.96 | 0.97 |
| 2gzaC | 0.66 | 0.72 | 0.71 | 4nj5A | 0.87 | 0.85 | 0.69 |
| 2hjqA | 0.82 | 0.82 | 0.55 | 4opaB | 0.64 | 0.64 | 0.64 |
| 2hwjA | 0.68 | 0.69 | 0.84 | 4qkuB | 0.98 | 1 | 0.96 |
| 2ijd1 | 0.74 | 0.74 | 0.74 | 4up9A | 0.95 | 0.97 | 0.72 |
| 2iu7A | 0.93 | 0.95 | 0.7 | 4w7sA | 0.71 | 0.63 | 0.75 |
| 2iw2A | 0.99 | 0.97 | 0.81 | 1bf2A | 0.98 | 0.97 | 0.87 |

| PDB ID | TM-score | | | PDB ID | TM-score | | |
|---|---|---|---|---|---|---|---|
| | E2EDA | E2EDA -w/o-S | E2EDA -w/o-S&T | | E2EDA | E2EDA -w/o-S | E2EDA -w/o-S&T |
| 2jz4A | 0.55 | 0.55 | 0.53 | 1bhgA | 0.96 | 0.82 | 0.82 |
| 2kdyA | 0.96 | 0.71 | 0.79 | 1f7uA | 0.97 | 0.91 | 0.86 |
| 2kn4A | 0.61 | 0.61 | 0.64 | 1fx7A | 0.66 | 0.66 | 0.62 |
| 2mbgA | 0.78 | 0.78 | 0.8 | 1griA | 0.72 | 0.59 | 0.63 |
| 2nsfA | 0.89 | 0.91 | 0.88 | 1h88C | 0.72 | 0.72 | 0.74 |
| 2nykA | 0.95 | 0.9 | 0.87 | 1m8pB | 0.96 | 0.96 | 0.8 |
| 2o6yA | 0.96 | 1 | 0.93 | 1ni5A | 0.71 | 0.64 | 0.66 |
| 2owbA | 0.98 | 0.96 | 0.96 | 1q25A | 0.7 | 0.69 | 0.73 |
| 2qfiA | 0.99 | 0.76 | 0.73 | 1uzjA | 0.77 | 0.74 | 0.72 |
| 2qp2A | 0.97 | 0.95 | 0.77 | 1zpuA | 0.89 | 0.89 | 0.77 |
| 2qygA | 0.99 | 1 | 0.94 | 1zy9A | 0.91 | 0.85 | 0.71 |
| 2r5wB | 0.97 | 0.93 | 0.63 | 2b5uA | 0.79 | 0.79 | 0.73 |
| 2uu7A | 0.98 | 0.91 | 0.95 | 2ewfA | 0.5 | 0.49 | 0.52 |
| 2w4bA | 0.98 | 0.99 | 0.92 | 2piaA | 0.72 | 0.71 | 0.67 |
| 2x7iA | 0.89 | 0.85 | 0.81 | 2r7dA | 0.98 | 0.95 | 0.82 |
| 2x8kC | 0.92 | 0.87 | 0.91 | 2uwnA | 0.8 | 0.75 | 0.63 |
| 2yilA | 0.62 | 0.63 | 0.64 | 2v0nA | 0.41 | 0.43 | 0.43 |
| 2yrqA | 0.52 | 0.52 | 0.65 | 2vgmA | 0.67 | 0.63 | 0.66 |
| 2zxcA | 0.99 | 1 | 0.87 | 2wqrB | 0.73 | 0.63 | 0.67 |
| 3a1iA | 0.9 | 0.9 | 0.9 | 2y25B | 0.63 | 0.63 | 0.63 |
| 3a45A | 0.98 | 0.97 | 0.92 | 2yk0A | 0.7 | 0.68 | 0.54 |
| 3a56A | 0.61 | 0.62 | 0.61 | 2zzqA | 0.97 | 0.93 | 0.43 |
| 3ajvA | 0.85 | 0.85 | 0.82 | 3bt1U | 0.89 | 0.86 | 0.55 |
| 3aqkA | 0.95 | 0.91 | 0.88 | 3c1yA | 0.99 | 0.98 | 0.82 |
| 3arbA | 0.96 | 0.94 | 0.93 | 3cw2C | 0.58 | 0.66 | 0.64 |
| 3aujG | 0.98 | 0.96 | 0.79 | 3f83A | 0.97 | 0.94 | 0.87 |
| 3b2zF | 0.95 | 0.98 | 0.88 | 3fc3A | 0.44 | 0.44 | 0.44 |
| 3b7wA | 0.96 | 0.83 | 0.82 | 3gbgA | 0.98 | 0.98 | 0.75 |
| 3bt3A | 0.85 | 0.93 | 0.81 | 3h5cB | 0.91 | 0.84 | 0.83 |
| 3c4tA | 0.83 | 0.74 | 0.85 | 3ibjA | 0.5 | 0.51 | 0.5 |
| 3craA | 0.93 | 0.97 | 0.91 | 3ippB | 0.94 | 0.91 | 0.74 |
| 3d30A | 0.89 | 0.91 | 0.89 | 3jymB | 0.92 | 0.92 | 0.79 |
| 3eo5A | 0.97 | 0.88 | 0.83 | 3kbgA | 0.86 | 0.56 | 0.46 |
| 3errA | 0.8 | 0.71 | 0.72 | 3mc8A | 0.68 | 0.67 | 0.59 |
| 3g79A | 0.95 | 0.97 | 0.99 | 3npfA | 0.84 | 0.72 | 0.64 |
| 3h2tA | 0.98 | 0.92 | 0.92 | 3orjA | 0.76 | 0.69 | 0.63 |
| 3hcsA | 0.82 | 0.71 | 0.73 | 3plaA | 0.66 | 0.71 | 0.71 |
| 3hyiA | 0.69 | 0.69 | 0.69 | 3qe9Y | 0.97 | 0.98 | 0.92 |
| 3i2dA | 0.87 | 0.96 | 0.79 | 3qjoA | 0.79 | 0.81 | 0.82 |
| 3iam2 | 0.99 | 0.94 | 0.8 | 3qphA | 0.71 | 0.7 | 0.54 |
| 3ifrA | 0.96 | 0.94 | 0.78 | 3qyeA | 0.97 | 0.92 | 0.96 |
| 3isqA | 0.95 | 0.95 | 0.76 | 3rimA | 0.96 | 0.97 | 0.92 |
| 3isqA | 0.94 | 0.91 | 0.79 | 3rrpA | 0.95 | 0.94 | 0.94 |
| 3k1rA | 0.62 | 0.63 | 0.62 | 3soaA | 0.67 | 0.68 | 0.68 |
| 3k2iA | 0.93 | 0.94 | 0.79 | 3tixD | 0.94 | 0.95 | 0.92 |
| 3kh5A | 0.95 | 0.94 | 0.87 | 3tp9A | 0.55 | 0.57 | 0.57 |
| 3kjpA | 0.86 | 0.52 | 0.8 | 3ua3A | 0.57 | 0.58 | 0.52 |
| 3kt1A | 0.97 | 0.91 | 0.58 | 3uj0A | 0.95 | 0.99 | 0.85 |
| 3ktmE | 0.73 | 0.61 | 0.76 | 3vn4A | 0.97 | 0.92 | 0.72 |
| 3kzwA | 1 | 0.95 | 0.97 | 3vsmA | 0.98 | 0.96 | 0.66 |
| 3l76A | 0.97 | 0.97 | 0.63 | 3w1bA | 0.77 | 0.76 | 0.77 |

| PDB ID | TM-score | | | PDB ID | TM-score | | |
|---|---|---|---|---|---|---|---|
| | E2EDA | E2EDA -w/o-S | E2EDA -w/o-S&T | | E2EDA | E2EDA -w/o-S | E2EDA -w/o-S&T |
| 3ld1A | 0.7 | 0.7 | 0.84 | 3zh9B | 0.72 | 0.64 | 0.62 |
| 3lsgA | 0.99 | 0.97 | 0.6 | 4alzA | 0.6 | 0.58 | 0.64 |
| 3me4A | 0.96 | 0.94 | 1 | 4ax8A | 0.69 | 0.69 | 0.63 |
| 3ml4C | 0.58 | 0.56 | 0.54 | 4b3iA | 0.85 | 0.82 | 0.83 |
| 3mx2B | 0.97 | 0.96 | 0.82 | 4bd9B | 0.67 | 0.6 | 0.6 |
| 3mzfA | 0.91 | 0.94 | 0.83 | 4c0aB | 0.92 | 0.92 | 0.69 |
| 3njaB | 0.59 | 0.63 | 0.6 | 4c0sA | 0.61 | 0.61 | 0.58 |
| 3nqiA | 0.93 | 0.93 | 0.75 | 4dimA | 0.76 | 0.84 | 0.75 |
| 3nt8A | 0.94 | 0.94 | 0.84 | 4indA | 0.45 | 0.43 | 0.53 |
| 3og5A | 0.9 | 0.97 | 0.96 | 4jdzB | 0.93 | 0.77 | 0.69 |
| 3oh0A | 0.94 | 0.89 | 0.87 | 4kc3B | 0.66 | 0.59 | 0.58 |
| 3pcsB | 0.99 | 1 | 0.99 | 4kikB | 0.82 | 0.68 | 0.64 |
| 3po3S | 0.93 | 0.93 | 0.85 | 4lmfA | 0.75 | 0.67 | 0.57 |
| 3pxpA | 0.74 | 0.69 | 0.7 | 4lziA | 0.47 | 0.47 | 0.64 |
| 3qavA | 0.94 | 0.95 | 0.99 | 4m9pA | 0.95 | 0.94 | 0.66 |
| 3qf4B | 0.9 | 0.92 | 0.81 | 4pt5A | 0.91 | 0.93 | 0.81 |
| 3qjjA | 0.93 | 0.95 | 0.87 | 4uwhA | 0.95 | 0.93 | 0.71 |
| 3qtdA | 0.98 | 0.9 | 0.88 | 1c1zA | 0.75 | 0.42 | 0.42 |
| 3r6bA | 0.97 | 0.84 | 0.84 | 1d2pA | 0.3 | 0.36 | 0.3 |
| 3rh7A | 0.99 | 0.99 | 0.71 | 1k7tA | 0.87 | 0.9 | 0.96 |
| 1kfqA | 0.92 | 0.88 | 0.91 | 2bydA | 0.85 | 0.89 | 0.89 |
| 1ldjA | 0.76 | 0.79 | 0.78 | 2c43A | 0.96 | 0.96 | 0.89 |
| 1nyqB | 0.91 | 0.97 | 0.89 | 2dfyC | 0.96 | 0.92 | 0.65 |
| 1ug9A | 0.64 | 0.66 | 0.69 | 2dlaA | 0.97 | 0.91 | 0.98 |
| 1z1wA | 0.96 | 0.94 | 0.98 | 2g3pA | 0.97 | 0.96 | 0.79 |
| 2au3A | 0.71 | 0.72 | 0.43 | 2gg6A | 1 | 0.97 | 0.98 |
| 2ii2A | 0.75 | 0.71 | 0.75 | 2gsyE | 0.95 | 0.94 | 0.88 |
| 2olsA | 0.46 | 0.45 | 0.48 | 2gzoA | 0.71 | 0.61 | 0.55 |
| 2ra1A | 0.6 | 0.44 | 0.44 | 2j2cA | 0.95 | 0.93 | 0.91 |
| 2v5dA | 0.75 | 0.81 | 0.74 | 2kfwA | 0.89 | 0.9 | 0.87 |
| 2xt6A | 0.98 | 0.97 | 0.98 | 2l9yA | 0.74 | 0.74 | 0.78 |
| 2zpaB | 0.94 | 0.8 | 0.59 | 2ntyB | 0.98 | 0.93 | 0.79 |
| 3apoA | 0.7 | 0.72 | 0.29 | 2r3vA | 0.97 | 0.98 | 0.89 |
| 3b43A | 0.36 | 0.49 | 0.36 | 2r58A | 0.99 | 0.96 | 0.8 |
| 3gf5B | 0.53 | 0.69 | 0.85 | 2w4mA | 0.96 | 0.95 | 0.76 |
| 3hjlA | 0.36 | 0.35 | 0.36 | 2x0cA | 0.67 | 0.71 | 0.63 |
| 3kq4B | 0.39 | 0.4 | 0.31 | 2y51A | 0.98 | 0.96 | 0.88 |
| 3kwlA | 0.47 | 0.41 | 0.36 | 2yb0E | 0.86 | 0.86 | 0.57 |
| 3ob8A | 0.95 | 0.93 | 0.91 | 2z86C | 0.96 | 0.9 | 0.86 |
| 3opfB | 0.77 | 0.56 | 0.62 | 3afoA | 0.91 | 0.76 | 0.59 |
| 3p53A | 0.46 | 0.47 | 0.74 | 3bu2A | 0.97 | 0.97 | 0.97 |
| 3pvlA | 0.9 | 0.79 | 0.73 | 3cvzA | 0.98 | 0.97 | 0.95 |
| 3r05A | 0.35 | 0.3 | 0.38 | 3dupA | 0.95 | 0.94 | 0.93 |
| 3ubhA | 0.65 | 0.83 | 0.65 | 3eswA | 0.95 | 0.94 | 0.94 |
| 3w2wA | 0.82 | 0.84 | 0.55 | 3eukH | 0.7 | 0.71 | 0.7 |
| 3zniA | 0.79 | 0.78 | 0.37 | 3fi7A | 0.95 | 0.9 | 0.93 |
| 4aimA | 0.8 | 0.89 | 0.84 | 3fvvA | 0.78 | 0.81 | 0.78 |
| 4ak1A | 0.28 | 0.3 | 0.3 | 3gmsA | 0.96 | 0.97 | 0.97 |
| 4aq1A | 0.81 | 0.52 | 0.28 | 3hzzB | 0.99 | 0.98 | 0.91 |
| 4fe9A | 0.91 | 0.76 | 0.31 | 3m1uA | 0.95 | 0.9 | 0.58 |
| 4h2aA | 0.77 | 0.75 | 0.82 | 3mw8A | 0.83 | 0.78 | 0.81 |
| 4i5sB | 0.5 | 0.42 | 0.57 | 3mwcA | 0.97 | 0.9 | 0.91 |

| PDB ID | TM-score | | | PDB ID | TM-score | | |
|---|---|---|---|---|---|---|---|
| | E2EDA | E2EDA -w/o-S | E2EDA -w/o-S&T | | E2EDA | E2EDA -w/o-S | E2EDA -w/o-S&T |
| 4iggB | 0.44 | 0.5 | 0.45 | 3nsjA | 0.97 | 0.94 | 0.97 |
| 4j9vA | 0.96 | 0.97 | 0.69 | 3ntkA | 0.95 | 0.97 | 0.94 |
| 4k3bA | 0.6 | 0.53 | 0.56 | 3oaaG | 0.95 | 0.93 | 0.86 |
| 4kwuA | 0.96 | 0.96 | 0.64 | 3ptyA | 0.93 | 0.93 | 0.92 |
| 4m00A | 0.49 | 0.5 | 0.53 | 3rfyA | 0.81 | 0.77 | 0.8 |
| 1bp1A | 0.98 | 0.95 | 0.96 | 3seoB | 0.66 | 0.62 | 0.62 |
| 1cjsA | 0.94 | 0.85 | 0.89 | 3spgA | 0.94 | 0.95 | 0.94 |
| 1ck1A | 0.99 | 0.97 | 0.97 | 3u0kA | 0.85 | 0.65 | 0.63 |
| 1ecrA | 1 | 0.99 | 0.99 | 3vlaA | 0.89 | 0.89 | 0.88 |
| 1f5qD | 0.91 | 0.92 | 0.84 | 3vstA | 0.96 | 0.99 | 0.87 |
| 1fa9A | 0.99 | 0.98 | 0.83 | 4aqfB | 0.83 | 0.86 | 0.77 |
| 1gu7A | 0.96 | 0.99 | 0.93 | 4b21A | 0.95 | 0.96 | 0.97 |
| 1itwA | 0.97 | 0.95 | 0.79 | 4dt4A | 0.84 | 0.77 | 0.91 |
| 1jkiA | 0.98 | 0.97 | 0.81 | 4dtfA | 0.84 | 0.95 | 0.85 |
| 1n80A | 0.96 | 0.98 | 0.95 | 4ewtA | 0.97 | 0.95 | 0.88 |
| 1nzjA | 0.97 | 0.97 | 0.71 | 4f23A | 0.92 | 0.85 | 0.85 |
| 1qhdA | 0.96 | 0.98 | 0.87 | 4fzbC | 0.86 | 0.93 | 0.74 |
| 1qz9A | 0.82 | 0.82 | 0.76 | 4g1pA | 0.96 | 0.93 | 0.93 |
| 1sb7B | 0.98 | 0.94 | 0.92 | 4gfqA | 0.8 | 0.86 | 0.8 |
| 1vk1A | 0.85 | 0.85 | 0.72 | 4hvzA | 0.98 | 0.92 | 0.9 |
| 1vrmA | 0.94 | 0.72 | 0.72 | 4il6B | 0.77 | 0.78 | 0.96 |
| 1xvuA | 0.97 | 0.96 | 0.98 | 4jxkA | 0.97 | 0.96 | 0.96 |
| 1yy3A | 0.91 | 0.78 | 0.81 | 4m8mB | 0.85 | 0.93 | 0.8 |
| 1z87A | 0.54 | 0.54 | 0.79 | 4mzyA | 0.97 | 0.96 | 0.97 |
| 2a1sC | 0.92 | 0.92 | 0.91 | 4onyA | 0.95 | 0.98 | 0.87 |
| 2a3lA | 0.97 | 0.98 | 0.98 | 4pyhA | 0.96 | 0.95 | 0.74 |
| 2bt1A | 0.97 | 0.96 | 0.92 | 4rg1A | 0.85 | 0.86 | 0.84 |

**Table S10.** Assembly results of E2EDA and E2EDA-w/o-R on benchmark proteins. #TM-score≥0.5 indicates the number of models with TM-score≥0.5. *P*-value is the result of Wilcoxon signed-rank test based on comparison with TM-scores of E2EDA.

| Method | TM-score | | | | | #TM-score ≥0.5 | *P*-value |
|---|---|---|---|---|---|---|---|
| | 2dis | 2dom | 3dom | m4dom | Total | | |
| **E2EDA** | **0.91** | **0.86** | **0.79** | **0.68** | **0.84** | **340** | NA |
| E2EDA-w/o-R | 0.79 | 0.78 | 0.65 | 0.54 | 0.72 | 316 | 7.3E-42 |

**Table S11.** Domain assembly results of E2EDA and E2EDA-w/o-R on 356 benchmark proteins.

| PDB ID | TM-score | | PDB ID | TM-score | | PDB ID | TM-score | |
|---|---|---|---|---|---|---|---|---|
| | E2EDA | E2EDA-w/o-R | | E2EDA | E2EDA-w/o-R | | E2EDA | E2EDA-w/o-R |
| 1cjyA | 0.82 | 0.83 | 3rwxA | 0.82 | 0.55 | 1kfqA | 0.92 | 0.61 |
| 1efdN | 0.92 | 0.84 | 3sb4A | 0.99 | 0.99 | 1ldjA | 0.76 | 0.46 |
| 1fjrA | 0.67 | 0.67 | 3swjA | 0.92 | 0.74 | 1nyqB | 0.91 | 0.84 |
| 1g87B | 0.98 | 0.86 | 3t58B | 0.94 | 0.53 | 1ug9A | 0.64 | 0.67 |
| 1hx6B | 0.8 | 0.61 | 3t7jA | 0.91 | 0.56 | 1z1wA | 0.96 | 0.92 |
| 1iwaA | 0.99 | 0.98 | 3u07C | 0.92 | 0.81 | 2au3A | 0.71 | 0.4 |
| 1m5qH | 0.59 | 0.57 | 3u0oB | 0.94 | 0.74 | 2ii2A | 0.75 | 0.79 |
| 1mkfA | 0.54 | 0.53 | 3u9gA | 0.75 | 0.75 | 2olsA | 0.46 | 0.45 |
| 1mkmB | 0.97 | 0.94 | 3ub1D | 0.99 | 0.99 | 2ra1A | 0.6 | 0.3 |
| 1nh2D | 0.92 | 0.95 | 3uitD | 0.55 | 0.55 | 2v5dA | 0.75 | 0.64 |
| 1pprM | 0.86 | 0.87 | 3uo3A | 0.87 | 0.53 | 2xt6A | 0.98 | 0.64 |
| 1prrA | 0.53 | 0.53 | 3v7oB | 0.63 | 0.64 | 2zpaB | 0.94 | 0.69 |
| 1q19A | 0.8 | 0.9 | 3vr8B | 0.95 | 0.92 | 3apoA | 0.7 | 0.3 |
| 1qwrA | 0.96 | 0.85 | 3wkuA | 0.77 | 0.77 | 3b43A | 0.36 | 0.25 |
| 1r71B | 0.84 | 0.53 | 3zvmA | 0.52 | 0.55 | 3gf5B | 0.53 | 0.3 |
| 1rh1A | 0.71 | 0.56 | 4acoA | 0.99 | 0.99 | 3hjlA | 0.36 | 0.36 |
| 1rktA | 0.9 | 0.9 | 4ap5A | 0.94 | 0.59 | 3kq4B | 0.39 | 0.34 |
| 1s6lA | 0.8 | 0.73 | 4axdA | 0.95 | 0.74 | 3kwlA | 0.47 | 0.43 |
| 1sp3A | 0.7 | 0.59 | 4bfiB | 0.85 | 0.85 | 3ob8A | 0.95 | 0.83 |
| 1vz6A | 0.98 | 0.95 | 4bt9B | 0.73 | 0.6 | 3opfB | 0.77 | 0.67 |
| 1w3aA | 0.59 | 0.57 | 4cczA | 0.87 | 0.63 | 3p53A | 0.46 | 0.46 |
| 1wv3A | 0.83 | 0.63 | 4d0nB | 0.89 | 0.67 | 3pvlA | 0.9 | 0.78 |
| 1x7pA | 0.94 | 0.65 | 4d1iG | 0.98 | 0.97 | 3r05A | 0.35 | 0.34 |
| 1x9yA | 0.84 | 0.56 | 4dj3A | 0.97 | 0.55 | 3ubhA | 0.65 | 0.54 |
| 1y11A | 0.61 | 0.62 | 4dqaA | 0.84 | 0.64 | 3w2wA | 0.82 | 0.72 |
| 1yiqA | 0.99 | 0.87 | 4eo3A | 0.57 | 0.64 | 3zniA | 0.79 | 0.79 |
| 1zbuB | 0.96 | 0.75 | 4eogA | 0.99 | 0.67 | 4aimA | 0.8 | 0.49 |
| 1ze1A | 0.94 | 0.92 | 4etxA | 0.98 | 0.96 | 4ak1A | 0.28 | 0.22 |
| 2ablA | 0.63 | 0.62 | 4fguA | 0.97 | 0.9 | 4aq1A | 0.81 | 0.47 |
| 2ahvA | 0.97 | 0.71 | 4fkcA | 0.95 | 0.62 | 4fe9A | 0.91 | 0.93 |
| 2bkpA | 0.6 | 0.61 | 4fxkC | 0.65 | 0.58 | 4h2aA | 0.77 | 0.61 |
| 2c1yA | 0.7 | 0.6 | 4gbyA | 0.99 | 0.99 | 4i5sB | 0.5 | 0.43 |
| 2cxcA | 0.89 | 0.88 | 4ggmX | 0.98 | 0.73 | 4iggB | 0.44 | 0.49 |
| 2d1cA | 0.94 | 0.79 | 4gslA | 0.96 | 0.96 | 4j9vA | 0.96 | 0.54 |
| 2d7iA | 0.77 | 0.77 | 4gyjA | 0.96 | 0.94 | 4k3bA | 0.6 | 0.51 |
| 2e9hA | 0.92 | 0.92 | 4h3tA | 0.92 | 0.96 | 4kwuA | 0.96 | 0.61 |
| 2e9xB | 0.9 | 0.63 | 4hmoA | 0.9 | 0.58 | 4m00A | 0.49 | 0.49 |
| 2evrA | 0.9 | 0.63 | 4ie6A | 0.99 | 0.69 | 1bp1A | 0.98 | 0.68 |
| 2ew9A | 0.56 | 0.57 | 4l5gA | 0.84 | 0.64 | 1cjsA | 0.94 | 0.58 |
| 2fd5A | 0.97 | 0.92 | 4lpqA | 0.95 | 0.95 | 1ck1A | 0.99 | 0.99 |
| 2gh8A | 0.9 | 0.65 | 4m8rA | 0.76 | 0.77 | 1ecrA | 1 | 1 |
| 2gt1A | 0.95 | 0.91 | 4n06B | 0.95 | 0.95 | 1f5qD | 0.91 | 0.58 |
| 2gzaC | 0.66 | 0.66 | 4nj5A | 0.87 | 0.6 | 1fa9A | 0.99 | 0.97 |
| 2hjqA | 0.82 | 0.53 | 4opaB | 0.64 | 0.71 | 1gu7A | 0.96 | 0.99 |
| 2hwjA | 0.68 | 0.67 | 4qkuB | 0.98 | 0.98 | 1itwA | 0.97 | 0.62 |
| 2ijd1 | 0.74 | 0.76 | 4up9A | 0.95 | 0.99 | 1jkiA | 0.98 | 0.97 |
| 2iu7A | 0.93 | 0.56 | 4w7sA | 0.71 | 0.71 | 1n80A | 0.96 | 0.96 |
| 2iw2A | 0.99 | 0.98 | 1bf2A | 0.98 | 0.86 | 1nzjA | 0.97 | 0.88 |

| PDB ID | TM-score | | PDB ID | TM-score | | PDB ID | TM-score | |
|---|---|---|---|---|---|---|---|---|
| | E2EDA | E2EDA-w/o-R | | E2EDA | E2EDA-w/o-R | | E2EDA | E2EDA-w/o-R |
| 2jz4A | 0.55 | 0.53 | 1bhgA | 0.96 | 0.86 | 1qhdA | 0.96 | 0.99 |
| 2kdyA | 0.96 | 0.96 | 1f7uA | 0.97 | 0.96 | 1qz9A | 0.82 | 0.66 |
| 2kn4A | 0.61 | 0.61 | 1fx7A | 0.66 | 0.71 | 1sb7B | 0.98 | 0.98 |
| 2mbgA | 0.78 | 0.78 | 1griA | 0.72 | 0.7 | 1vk1A | 0.85 | 0.64 |
| 2nsfA | 0.89 | 0.81 | 1h88C | 0.72 | 0.38 | 1vrmA | 0.94 | 0.76 |
| 2nykA | 0.95 | 0.95 | 1m8pB | 0.96 | 0.86 | 1xvuA | 0.97 | 0.77 |
| 2o6yA | 0.96 | 0.94 | 1ni5A | 0.71 | 0.63 | 1yy3A | 0.91 | 0.88 |
| 2owbA | 0.98 | 0.98 | 1q25A | 0.7 | 0.69 | 1z87A | 0.54 | 0.52 |
| 2qfiA | 0.99 | 0.78 | 1uzjA | 0.77 | 0.74 | 2a1sC | 0.92 | 0.82 |
| 2qp2A | 0.97 | 0.97 | 1zpuA | 0.89 | 0.75 | 2a3lA | 0.97 | 0.97 |
| 2qygA | 0.99 | 0.96 | 1zy9A | 0.91 | 0.85 | 2bt1A | 0.97 | 0.86 |
| 2r5wB | 0.97 | 0.95 | 2b5uA | 0.79 | 0.47 | 2bydA | 0.85 | 0.59 |
| 2uu7A | 0.98 | 0.72 | 2ewfA | 0.5 | 0.48 | 2c43A | 0.96 | 0.56 |
| 2w4bA | 0.98 | 0.95 | 2piaA | 0.72 | 0.42 | 2dfyC | 0.96 | 0.52 |
| 2x7iA | 0.89 | 0.89 | 2r7dA | 0.98 | 0.86 | 2dlaA | 0.97 | 0.78 |
| 2x8kC | 0.92 | 0.9 | 2uwnA | 0.8 | 0.74 | 2g3pA | 0.97 | 0.93 |
| 2yilA | 0.62 | 0.62 | 2v0nA | 0.41 | 0.39 | 2gg6A | 1 | 1 |
| 2yrqA | 0.52 | 0.51 | 2vgmA | 0.67 | 0.47 | 2gsyE | 0.95 | 0.68 |
| 2zxcA | 0.99 | 0.99 | 2wqrB | 0.73 | 0.48 | 2gzoA | 0.71 | 0.56 |
| 3a1iA | 0.9 | 0.9 | 2y25B | 0.63 | 0.4 | 2j2cA | 0.95 | 0.69 |
| 3a45A | 0.98 | 0.98 | 2yk0A | 0.7 | 0.45 | 2kfwA | 0.89 | 0.75 |
| 3a56A | 0.61 | 0.61 | 2zzqA | 0.97 | 0.76 | 2l9yA | 0.74 | 0.74 |
| 3ajvA | 0.85 | 0.53 | 3bt1U | 0.89 | 0.65 | 2ntyB | 0.98 | 0.98 |
| 3aqkA | 0.95 | 0.88 | 3c1yA | 0.99 | 0.87 | 2r3vA | 0.97 | 0.92 |
| 3arbA | 0.96 | 0.68 | 3cw2C | 0.58 | 0.59 | 2r58A | 0.99 | 0.99 |
| 3aujG | 0.98 | 0.96 | 3f83A | 0.97 | 0.68 | 2w4mA | 0.96 | 0.96 |
| 3b2zF | 0.95 | 0.94 | 3fc3A | 0.44 | 0.43 | 2x0cA | 0.67 | 0.63 |
| 3b7wA | 0.96 | 0.8 | 3gbgA | 0.98 | 0.97 | 2y51A | 0.98 | 0.99 |
| 3bt3A | 0.85 | 0.89 | 3h5cB | 0.91 | 0.51 | 2yb0E | 0.86 | 0.85 |
| 3c4tA | 0.83 | 0.83 | 3ibjA | 0.5 | 0.49 | 2z86C | 0.96 | 0.6 |
| 3craA | 0.93 | 0.93 | 3ippB | 0.94 | 0.91 | 3afoA | 0.91 | 0.56 |
| 3d30A | 0.89 | 0.53 | 3jymB | 0.92 | 0.92 | 3bu2A | 0.97 | 0.64 |
| 3eo5A | 0.97 | 0.89 | 3kbgA | 0.86 | 0.9 | 3cvzA | 0.98 | 0.97 |
| 3errA | 0.8 | 0.62 | 3mc8A | 0.68 | 0.59 | 3dupA | 0.95 | 0.95 |
| 3g79A | 0.95 | 0.65 | 3npfA | 0.84 | 0.51 | 3eswA | 0.95 | 0.85 |
| 3h2tA | 0.98 | 0.92 | 3orjA | 0.76 | 0.72 | 3eukH | 0.7 | 0.75 |
| 3hcsA | 0.82 | 0.69 | 3plaA | 0.66 | 0.67 | 3fi7A | 0.95 | 0.67 |
| 3hyiA | 0.69 | 0.68 | 3qe9Y | 0.97 | 0.62 | 3fvvA | 0.78 | 0.78 |
| 3i2dA | 0.87 | 0.58 | 3qjoA | 0.79 | 0.78 | 3gmsA | 0.96 | 0.66 |
| 3iam2 | 0.99 | 0.99 | 3qphA | 0.71 | 0.69 | 3hzzB | 0.99 | 0.98 |
| 3ifrA | 0.96 | 0.92 | 3qyeA | 0.97 | 0.74 | 3m1uA | 0.95 | 0.95 |
| 3isqA | 0.95 | 0.95 | 3rimA | 0.96 | 0.77 | 3mw8A | 0.83 | 0.83 |
| 3isqA | 0.94 | 0.92 | 3rrpA | 0.95 | 0.63 | 3mwcA | 0.97 | 0.84 |
| 3k1rA | 0.62 | 0.62 | 3soaA | 0.67 | 0.68 | 3nsjA | 0.97 | 0.88 |
| 3k2iA | 0.93 | 0.84 | 3tixD | 0.94 | 0.61 | 3ntkA | 0.95 | 0.95 |
| 3kh5A | 0.95 | 0.97 | 3tp9A | 0.55 | 0.55 | 3oaaG | 0.95 | 0.73 |
| 3kjpA | 0.86 | 0.63 | 3ua3A | 0.57 | 0.45 | 3ptyA | 0.93 | 0.93 |
| 3kt1A | 0.97 | 0.98 | 3uj0A | 0.95 | 0.47 | 3rfyA | 0.81 | 0.72 |
| 3ktmE | 0.73 | 0.73 | 3vn4A | 0.97 | 0.88 | 3seoB | 0.66 | 0.65 |
| 3kzwA | 1 | 0.96 | 3vsmA | 0.98 | 0.97 | 3spgA | 0.94 | 0.62 |
| 3l76A | 0.97 | 0.95 | 3w1bA | 0.77 | 0.74 | 3u0kA | 0.85 | 0.85 |
| 3ld1A | 0.7 | 0.7 | 3zh9B | 0.72 | 0.56 | 3vlaA | 0.89 | 0.65 |

| PDB ID | TM-score | | PDB ID | TM-score | | PDB ID | TM-score | |
|---|---|---|---|---|---|---|---|---|
| | E2EDA | E2EDA-w/o-R | | E2EDA | E2EDA-w/o-R | | E2EDA | E2EDA-w/o-R |
| 3lsgA | 0.99 | 0.56 | 4alzA | 0.6 | 0.4 | 3vstA | 0.96 | 0.98 |
| 3me4A | 0.96 | 0.51 | 4ax8A | 0.69 | 0.52 | 4aqfB | 0.83 | 0.78 |
| 3ml4C | 0.58 | 0.98 | 4b3iA | 0.85 | 0.86 | 4b21A | 0.95 | 0.6 |
| 3mx2B | 0.97 | 0.97 | 4bd9B | 0.67 | 0.38 | 4dt4A | 0.84 | 0.85 |
| 3mzfA | 0.91 | 0.84 | 4c0aB | 0.92 | 0.66 | 4dtfA | 0.84 | 0.61 |
| 3njaB | 0.59 | 0.59 | 4c0sA | 0.61 | 0.55 | 4ewtA | 0.97 | 0.74 |
| 3nqiA | 0.93 | 0.75 | 4dimA | 0.76 | 0.81 | 4f23A | 0.92 | 0.9 |
| 3nt8A | 0.94 | 0.54 | 4indA | 0.45 | 0.41 | 4fzbC | 0.86 | 0.77 |
| 3og5A | 0.9 | 0.9 | 4jdzB | 0.93 | 0.55 | 4g1pA | 0.96 | 0.94 |
| 3oh0A | 0.94 | 0.94 | 4kc3B | 0.66 | 0.59 | 4gfqA | 0.8 | 0.61 |
| 3pcsB | 0.99 | 0.75 | 4kikB | 0.82 | 0.5 | 4hvzA | 0.98 | 0.98 |
| 3po3S | 0.93 | 0.56 | 4lmfA | 0.75 | 0.48 | 4il6B | 0.77 | 0.65 |
| 3pxpA | 0.74 | 0.7 | 4lziA | 0.47 | 0.47 | 4jxkA | 0.97 | 0.97 |
| 3qavA | 0.94 | 0.94 | 4m9pA | 0.95 | 0.95 | 4m8mB | 0.85 | 0.8 |
| 3qf4B | 0.9 | 0.59 | 4pt5A | 0.91 | 0.82 | 4mzyA | 0.97 | 0.71 |
| 3qjjA | 0.93 | 0.54 | 4uwhA | 0.95 | 0.46 | 4onyA | 0.95 | 0.58 |
| 3qtdA | 0.98 | 0.65 | 1c1zA | 0.75 | 0.48 | 4pyhA | 0.96 | 0.96 |
| 3r6bA | 0.97 | 0.91 | 1d2pA | 0.3 | 0.29 | 4rg1A | 0.85 | 0.6 |
| 3rh7A | 0.99 | 0.99 | 1k7tA | 0.87 | 0.48 | | | |

**Table S12.** Summary of the best model during E2EDA assembly and the final first model selected by RMscore on the benchmark set.

| Type | TM-score | | | | | *P*-value |
|---|---|---|---|---|---|---|
| | 2dis | 2dom | 3dom | m4dom | Total | |
| The best model during assembly | 0.92 | 0.88 | 0.80 | 0.69 | 0.85 | NA |
| Final first model | 0.91 | 0.86 | 0.79 | 0.68 | 0.84 | 1.8E-20 |

**Table S13.** Detailed results of the best model during E2EDA assembly and the final first model selected by RMscore on the benchmark set.

| PDB ID | TM-score | | PDB ID | TM-score | | PDB ID | TM-score | |
|---|---|---|---|---|---|---|---|---|
| | Final first model | Best model | | Final first model | Best model | | Final first model | Best model |
| 1cjyA | 0.82 | 0.88 | 3rwxA | 0.82 | 0.82 | 1kfqA | 0.92 | 0.94 |
| 1efdN | 0.92 | 0.92 | 3sb4A | 0.99 | 0.99 | 1ldjA | 0.76 | 0.76 |
| 1fjrA | 0.67 | 0.69 | 3swjA | 0.92 | 0.92 | 1nyqB | 0.91 | 0.91 |
| 1g87B | 0.98 | 0.98 | 3t58B | 0.94 | 0.95 | 1ug9A | 0.64 | 0.64 |
| 1hx6B | 0.8 | 0.83 | 3t7jA | 0.91 | 0.91 | 1z1wA | 0.96 | 0.96 |
| 1iwaA | 0.99 | 0.99 | 3u07C | 0.92 | 0.92 | 2au3A | 0.71 | 0.71 |
| 1m5qH | 0.59 | 0.6 | 3u0oB | 0.94 | 0.94 | 2ii2A | 0.75 | 0.79 |
| 1mkfA | 0.54 | 0.56 | 3u9gA | 0.75 | 0.79 | 2olsA | 0.46 | 0.51 |
| 1mkm | 0.97 | 0.97 | 3ub1D | 0.99 | 0.99 | 2ra1A | 0.6 | 0.6 |
| 1nh2D | 0.92 | 0.95 | 3uitD | 0.55 | 0.55 | 2v5dA | 0.75 | 0.76 |
| 1pprM | 0.86 | 0.98 | 3uo3A | 0.87 | 0.87 | 2xt6A | 0.98 | 0.98 |
| 1prrA | 0.53 | 0.62 | 3v7oB | 0.63 | 0.64 | 2zpaB | 0.94 | 0.94 |
| 1q19A | 0.8 | 0.9 | 3vr8B | 0.95 | 0.95 | 3apoA | 0.7 | 0.71 |
| 1qwrA | 0.96 | 0.96 | 3wkuA | 0.77 | 0.8 | 3b43A | 0.36 | 0.36 |
| 1r71B | 0.84 | 0.86 | 3zvmA | 0.52 | 0.56 | 3gf5B | 0.53 | 0.6 |
| 1rh1A | 0.71 | 0.87 | 4acoA | 0.99 | 0.99 | 3hjlA | 0.36 | 0.36 |
| 1rktA | 0.9 | 0.93 | 4ap5A | 0.94 | 0.97 | 3kq4B | 0.39 | 0.4 |
| 1s6lA | 0.8 | 0.8 | 4axdA | 0.95 | 0.95 | 3kwlA | 0.47 | 0.47 |
| 1sp3A | 0.7 | 0.7 | 4bfiB | 0.85 | 0.93 | 3ob8A | 0.95 | 0.95 |
| 1vz6A | 0.98 | 0.98 | 4bt9B | 0.73 | 0.84 | 3opfB | 0.77 | 0.79 |
| 1w3aA | 0.59 | 0.8 | 4cczA | 0.87 | 0.97 | 3p53A | 0.46 | 0.46 |
| 1wv3A | 0.83 | 0.85 | 4d0nB | 0.89 | 0.95 | 3pvlA | 0.9 | 0.92 |
| 1x7pA | 0.94 | 0.95 | 4d1iG | 0.98 | 0.98 | 3r05A | 0.35 | 0.35 |
| 1x9yA | 0.84 | 0.84 | 4dj3A | 0.97 | 0.97 | 3ubhA | 0.65 | 0.67 |
| 1y11A | 0.61 | 0.71 | 4dqaA | 0.84 | 0.84 | 3w2wA | 0.82 | 0.85 |
| 1yiqA | 0.99 | 0.99 | 4eo3A | 0.57 | 0.57 | 3zniA | 0.79 | 0.81 |
| 1zbuB | 0.96 | 0.96 | 4eogA | 0.99 | 0.99 | 4aimA | 0.8 | 0.83 |
| 1ze1A | 0.94 | 0.95 | 4etxA | 0.98 | 0.99 | 4ak1A | 0.28 | 0.32 |
| 2ablA | 0.63 | 0.64 | 4fguA | 0.97 | 0.97 | 4aq1A | 0.81 | 0.81 |
| 2ahvA | 0.97 | 0.97 | 4fkcA | 0.95 | 0.98 | 4fe9A | 0.91 | 0.91 |
| 2bkpA | 0.6 | 0.71 | 4fxkC | 0.65 | 0.65 | 4h2aA | 0.77 | 0.77 |
| 2c1yA | 0.7 | 0.7 | 4gbyA | 0.99 | 0.99 | 4i5sB | 0.5 | 0.56 |
| 2cxcA | 0.89 | 0.89 | 4ggmX | 0.98 | 0.98 | 4iggB | 0.44 | 0.44 |
| 2d1cA | 0.94 | 0.94 | 4gslA | 0.96 | 0.99 | 4j9vA | 0.96 | 0.96 |
| 2d7iA | 0.77 | 0.77 | 4gyjA | 0.96 | 0.96 | 4k3bA | 0.6 | 0.6 |
| 2e9hA | 0.92 | 0.92 | 4h3tA | 0.92 | 0.96 | 4kwuA | 0.96 | 0.96 |
| 2e9xB | 0.9 | 0.9 | 4hmoA | 0.9 | 0.9 | 4m00A | 0.49 | 0.49 |
| 2evrA | 0.9 | 0.91 | 4ie6A | 0.99 | 0.99 | 1bp1A | 0.98 | 0.98 |
| 2ew9A | 0.56 | 0.58 | 4l5gA | 0.84 | 0.91 | 1cjsA | 0.94 | 0.94 |
| 2fd5A | 0.97 | 0.97 | 4lpqA | 0.95 | 0.95 | 1ck1A | 0.99 | 1 |
| 2gh8A | 0.9 | 0.9 | 4m8rA | 0.76 | 0.77 | 1ecrA | 1 | 1 |
| 2gt1A | 0.95 | 0.98 | 4n06B | 0.95 | 0.95 | 1f5qD | 0.91 | 0.91 |
| 2gzaC | 0.66 | 0.68 | 4nj5A | 0.87 | 0.87 | 1fa9A | 0.99 | 1 |
| 2hjqA | 0.82 | 0.82 | 4opaB | 0.64 | 0.84 | 1gu7A | 0.96 | 0.97 |
| 2hwjA | 0.68 | 0.73 | 4qkuB | 0.98 | 0.98 | 1itwA | 0.97 | 0.97 |
| 2ijd1 | 0.74 | 0.79 | 4up9A | 0.95 | 0.99 | 1jkiA | 0.98 | 0.98 |
| 2iu7A | 0.93 | 0.96 | 4w7sA | 0.71 | 0.89 | 1n80A | 0.96 | 0.98 |
| 2iw2A | 0.99 | 0.99 | 1bf2A | 0.98 | 0.98 | 1nzjA | 0.97 | 0.98 |

| PDB ID | TM-score | | PDB ID | TM-score | | PDB ID | TM-score | |
|---|---|---|---|---|---|---|---|---|
| | Final first model | Best model | | Final first model | Best model | | Final first model | Best model |
| 2jz4A | 0.55 | 0.55 | 1bhgA | 0.96 | 0.96 | 1qhdA | 0.96 | 0.96 |
| 2kdyA | 0.96 | 0.96 | 1f7uA | 0.97 | 0.97 | 1qz9A | 0.82 | 0.82 |
| 2kn4A | 0.61 | 0.61 | 1fx7A | 0.66 | 0.68 | 1sb7B | 0.98 | 0.98 |
| 2mbgA | 0.78 | 0.79 | 1griA | 0.72 | 0.72 | 1vk1A | 0.85 | 0.85 |
| 2nsfA | 0.89 | 0.89 | 1h88C | 0.72 | 0.72 | 1vrmA | 0.94 | 0.94 |
| 2nykA | 0.95 | 0.95 | 1m8pB | 0.96 | 0.96 | 1xvuA | 0.97 | 0.98 |
| 2o6yA | 0.96 | 0.96 | 1ni5A | 0.71 | 0.84 | 1yy3A | 0.91 | 0.95 |
| 2owbA | 0.98 | 0.98 | 1q25A | 0.7 | 0.71 | 1z87A | 0.54 | 0.55 |
| 2qfiA | 0.99 | 0.99 | 1uzjA | 0.77 | 0.77 | 2a1sC | 0.92 | 0.95 |
| 2qp2A | 0.97 | 0.97 | 1zpuA | 0.89 | 0.89 | 2a3lA | 0.97 | 0.98 |
| 2qygA | 0.99 | 0.99 | 1zy9A | 0.91 | 0.91 | 2bt1A | 0.97 | 0.97 |
| 2r5wB | 0.97 | 0.97 | 2b5uA | 0.79 | 0.89 | 2bydA | 0.85 | 0.85 |
| 2uu7A | 0.98 | 0.98 | 2ewfA | 0.5 | 0.51 | 2c43A | 0.96 | 0.96 |
| 2w4bA | 0.98 | 0.98 | 2piaA | 0.72 | 0.75 | 2dfyC | 0.96 | 0.96 |
| 2x7iA | 0.89 | 0.91 | 2r7dA | 0.98 | 0.98 | 2dlaA | 0.97 | 0.99 |
| 2x8kC | 0.92 | 0.92 | 2uwnA | 0.8 | 0.83 | 2g3pA | 0.97 | 0.97 |
| 2yilA | 0.62 | 0.66 | 2v0nA | 0.41 | 0.41 | 2gg6A | 1 | 1 |
| 2yrqA | 0.52 | 0.52 | 2vgmA | 0.67 | 0.68 | 2gsyE | 0.95 | 0.95 |
| 2zxcA | 0.99 | 1 | 2wqrB | 0.73 | 0.73 | 2gzoA | 0.71 | 0.71 |
| 3a1iA | 0.9 | 0.9 | 2y25B | 0.63 | 0.63 | 2j2cA | 0.95 | 0.95 |
| 3a45A | 0.98 | 0.98 | 2yk0A | 0.7 | 0.71 | 2kfwA | 0.89 | 0.89 |
| 3a56A | 0.61 | 0.64 | 2zzqA | 0.97 | 0.99 | 2l9yA | 0.74 | 0.74 |
| 3ajvA | 0.85 | 0.86 | 3bt1U | 0.89 | 0.89 | 2ntyB | 0.98 | 0.99 |
| 3aqkA | 0.95 | 0.95 | 3c1yA | 0.99 | 0.99 | 2r3vA | 0.97 | 0.97 |
| 3arbA | 0.96 | 0.96 | 3cw2C | 0.58 | 0.64 | 2r58A | 0.99 | 0.99 |
| 3aujG | 0.98 | 0.98 | 3f83A | 0.97 | 0.97 | 2w4mA | 0.96 | 0.98 |
| 3b2zF | 0.95 | 0.95 | 3fc3A | 0.44 | 0.46 | 2x0cA | 0.67 | 0.69 |
| 3b7wA | 0.96 | 0.96 | 3gbgA | 0.98 | 0.98 | 2y51A | 0.98 | 0.99 |
| 3bt3A | 0.85 | 0.89 | 3h5cB | 0.91 | 0.91 | 2yb0E | 0.86 | 0.92 |
| 3c4tA | 0.83 | 0.89 | 3ibjA | 0.5 | 0.56 | 2z86C | 0.96 | 0.96 |
| 3craA | 0.93 | 0.93 | 3ippB | 0.94 | 0.94 | 3afoA | 0.91 | 0.91 |
| 3d30A | 0.89 | 0.89 | 3jymB | 0.92 | 0.96 | 3bu2A | 0.97 | 0.97 |
| 3eo5A | 0.97 | 0.97 | 3kbgA | 0.86 | 0.94 | 3cvzA | 0.98 | 0.98 |
| 3errA | 0.8 | 0.84 | 3mc8A | 0.68 | 0.71 | 3dupA | 0.95 | 0.95 |
| 3g79A | 0.95 | 0.95 | 3npfA | 0.84 | 0.84 | 3eswA | 0.95 | 0.98 |
| 3h2tA | 0.98 | 0.98 | 3orjA | 0.76 | 0.81 | 3eukH | 0.7 | 0.76 |
| 3hcsA | 0.82 | 0.84 | 3plaA | 0.66 | 0.66 | 3fi7A | 0.95 | 0.95 |
| 3hyiA | 0.69 | 0.7 | 3qe9Y | 0.97 | 0.97 | 3fvvA | 0.78 | 0.78 |
| 3i2dA | 0.87 | 0.9 | 3qjoA | 0.79 | 0.81 | 3gmsA | 0.96 | 0.96 |
| 3iam2 | 0.99 | 0.99 | 3qphA | 0.71 | 0.71 | 3hzzB | 0.99 | 0.99 |
| 3ifrA | 0.96 | 0.98 | 3qyeA | 0.97 | 0.97 | 3m1uA | 0.95 | 0.95 |
| 3isqA | 0.95 | 0.96 | 3rimA | 0.96 | 0.97 | 3mw8A | 0.83 | 0.86 |
| 3isqA | 0.94 | 0.94 | 3rrpA | 0.95 | 0.96 | 3mwcA | 0.97 | 0.97 |
| 3k1rA | 0.62 | 0.65 | 3soaA | 0.67 | 0.72 | 3nsjA | 0.97 | 0.98 |
| 3k2iA | 0.93 | 0.94 | 3tixD | 0.94 | 0.94 | 3ntkA | 0.95 | 0.95 |
| 3kh5A | 0.95 | 0.97 | 3tp9A | 0.55 | 0.57 | 3oaaG | 0.95 | 0.96 |
| 3kjpA | 0.86 | 0.86 | 3ua3A | 0.57 | 0.57 | 3ptyA | 0.93 | 0.97 |
| 3kt1A | 0.97 | 0.98 | 3uj0A | 0.95 | 0.96 | 3rfyA | 0.81 | 0.82 |
| 3ktmE | 0.73 | 0.73 | 3vn4A | 0.97 | 0.97 | 3seoB | 0.66 | 0.66 |
| 3kzwA | 1 | 1 | 3vsmA | 0.98 | 0.98 | 3spgA | 0.94 | 0.95 |
| 3l76A | 0.97 | 0.97 | 3w1bA | 0.77 | 0.78 | 3u0kA | 0.85 | 0.85 |
| 3ld1A | 0.7 | 0.72 | 3zh9B | 0.72 | 0.72 | 3vlaA | 0.89 | 0.93 |

| PDB ID | TM-score | | PDB ID | TM-score | | PDB ID | TM-score | |
|---|---|---|---|---|---|---|---|---|
| | Final first model | Best model | | Final first model | Best model | | Final first model | Best model |
| 3lsgA | 0.99 | 0.99 | 4alzA | 0.6 | 0.62 | 3vstA | 0.96 | 0.98 |
| 3me4A | 0.96 | 0.96 | 4ax8A | 0.69 | 0.69 | 4aqfB | 0.83 | 0.92 |
| 3ml4C | 0.58 | 0.74 | 4b3iA | 0.85 | 0.91 | 4b21A | 0.95 | 0.95 |
| 3mx2B | 0.97 | 0.97 | 4bd9B | 0.67 | 0.7 | 4dt4A | 0.84 | 0.85 |
| 3mzfA | 0.91 | 0.95 | 4c0aB | 0.92 | 0.92 | 4dtfA | 0.84 | 0.84 |
| 3njaB | 0.59 | 0.61 | 4c0sA | 0.61 | 0.61 | 4ewtA | 0.97 | 0.98 |
| 3nqiA | 0.93 | 0.93 | 4dimA | 0.76 | 0.86 | 4f23A | 0.92 | 0.92 |
| 3nt8A | 0.94 | 0.94 | 4indA | 0.45 | 0.46 | 4fzbC | 0.86 | 0.86 |
| 3og5A | 0.9 | 0.9 | 4jdzB | 0.93 | 0.98 | 4g1pA | 0.96 | 0.97 |
| 3oh0A | 0.94 | 0.94 | 4kc3B | 0.66 | 0.66 | 4gfqA | 0.8 | 0.86 |
| 3pcsB | 0.99 | 1 | 4kikB | 0.82 | 0.82 | 4hvzA | 0.98 | 0.98 |
| 3po3S | 0.93 | 0.93 | 4lmfA | 0.75 | 0.75 | 4il6B | 0.77 | 0.94 |
| 3pxpA | 0.74 | 0.74 | 4lziA | 0.47 | 0.54 | 4jxkA | 0.97 | 0.99 |
| 3qavA | 0.94 | 0.94 | 4m9pA | 0.95 | 0.95 | 4m8mB | 0.85 | 0.85 |
| 3qf4B | 0.9 | 0.93 | 4pt5A | 0.91 | 0.93 | 4mzyA | 0.97 | 0.97 |
| 3qjjA | 0.93 | 0.93 | 4uwhA | 0.95 | 0.95 | 4onyA | 0.95 | 0.95 |
| 3qtdA | 0.98 | 0.98 | 1c1zA | 0.75 | 0.75 | 4pyhA | 0.96 | 0.96 |
| 3r6bA | 0.97 | 0.97 | 1d2pA | 0.3 | 0.33 | 4rg1A | 0.85 | 0.86 |
| 3rh7A | 0.99 | 0.99 | 1k7tA | 0.87 | 0.87 | | | |
